# MolX: A Geometric Foundation Model for Protein-Ligand Modelling

**DOI:** 10.64898/2026.02.26.708362

**Authors:** Jie Liu, Tong Pan, Xudong Guo, Zixu Ran, Yi Hao, Yannan Yang, Ashley P. Ng, Shirui Pan, Jiangning Song, Fuyi Li

## Abstract

Understanding how small molecules interact with protein binding pockets is central to structure-based drug discovery. Accurately modelling these interactions requires capturing the 3D geometry and physicochemical complementarity of binding interfaces, yet existing computational approaches encode proteins and ligands separately or rely on simplified structural representations that do not explicitly model cross-entity spatial relationships. Such decoupled representations restrict their capacity to capture interface-level geometric constraints that arise from protein-ligand coorganisation. Here we present MolX, a Graph Transformer foundation model that jointly learns geometric and chemical representations of protein pockets and ligands from large-scale 3D structural data. Integrating over 3 million protein pockets and 5 million molecules, MolX represents both entities as E(3)-equivariant graphs to preserve spatial geometry and chemical context. The architecture employs dual E(3)-equivariant graph Transformer encoders to model pocket and ligand embeddings, ensuring representations remain invariant to rotation, translation, and reflection. MolX is pretrained using a hybrid learning paradigm that combines supervised biochemical objectives, logP and energy-gap regression, with self-supervised geometric objectives, coordinate reconstruction, and atom-type prediction, fostering generalisable molecular understanding. Across eight downstream benchmarks, including antibody-drug conjugates (ADC), proteolysis-targeting chimeras (PROTAC), molecular glue, and PCBA activity prediction, as well as binding affinity and physicochemical property regression, MolX achieves consistent state-of-the-art performance and strong cross-domain generalisation. Furthermore, MolX incorporates a sparse autoencoder module to decompose latent representations into interpretable biological components, thereby revealing the pocket-ligand interactions that drive prediction outcomes. Together, MolX establishes a scalable and interpretable foundation model for molecular representation learning, providing a unified framework for predicting and interpreting complex small-molecule-protein interactions.

## Introduction

Deciphering how molecular structures encode biological function remains one of the central challenges in modern biochemistry and drug discovery [1, 2]. Despite decades of progress in structural biology and computational chemistry, our ability to accurately predict how small molecules bind to protein pockets remains limited. This challenge arises from the complex interplay of three-dimensional geometry, shape complementarity, and physicochemical interactions that determine binding affinity and selectivity [3, 4]. Bridging this gap requires artificial intelligence models that capture these structural principles in a unified, biologically meaningful way [5, 6].

Existing computational strategies for modelling protein-ligand interactions can be broadly categorised as sequence-based and structure-based approaches according to the type of input they require [5, 7]. Sequence-based methods operate on one-dimensional representations, such as ligand molecular strings (SMILES) and protein amino-acid sequences [8, 9, 10]. Despite their simplicity and scalability in input, sequence-based models omit explicit three-dimensional structural information, limiting their ability to capture interaction geometry and physicochemical constraints [11, 12, 13, 14]. Structure-based methods overcome this limitation by incorporating three-dimensional protein and ligand structures to explicitly model spatial complementarity [15, 16], which is difficult to infer from sequence alone. Despite their effectiveness, a large fraction of current 3D models treat proteins and ligands independently or focus only on local atomic geometry [17, 18, 19, 20], thereby missing the interaction-specific patterns that emerge when both components are jointly represented.

Graph-based learning strategies provide a natural way to effectively model three-dimensional structural information [21, 22]. Yet the standard Transformer architecture, as a common backbone, was originally developed for sequential data, where structural information can be incorporated by assigning an embedding to each position [23, 19]. However, molecular and protein structures are inherently non-sequential [24]. Atoms and residues are embedded in a three-dimensional space and connected through complex topological relationships rather than arranged along a linear order [25, 26, 27]. As a result, positional encoding strategies designed for sequences are insufficient to capture the geometric and relational structure of molecular graphs [28, 29, 30]. Therefore, effectively modelling protein pockets and small molecules requires architectures that can encode spatial relationships and graph connectivity in a manner consistent with three-dimensional geometry [31, 32].

Building on these considerations, we introduce MolX, a pretrained foundation model that jointly learns protein pocket and small-molecule ligand representations from large-scale 3D structural data. By representing both entities as 3D graphs and encoding them with E(3)-equivariant graph Transformer layers [19, 33], MolX captures geometry-aware interaction patterns that are invariant to global spatial transformations. The model is pretrained using a hybrid strategy that inherently combines supervised prediction of physicochemical properties with self-supervised reconstruction of corrupted atomic coordinates and atom identities, thereby encouraging the learning of transferable, structure-informed representations [34]. Consequently, MolX can be efficiently fine-tuned for a wide range of downstream protein-ligand tasks, consistently achieving state-of-the-art performance across diverse classification and regression benchmarks. Furthermore, MolX integrates a sparse autoencoder [35, 36] to decompose latent representations into interpretable features. This enables systematic attribution of model predictions to specific protein regions and molecular sub-structures, and provides mechanistic insights into learned interaction motifs.

## Results

### MolX Framework

MolX is a pretrained foundation model that jointly learns representations of ligands and protein pockets from large-scale three-dimensional structural data. After lightweight fine-tuning, MolX can be easily adapted to a wide range of downstream protein-ligand interaction prediction tasks (Fig. 1a). The inputs to MolX consist of protein pockets and ligand molecules (Fig. 1b). Both pockets and ligands are represented as three-dimensional graphs, where nodes correspond to atoms and edges represent chemical bonds. Each atom is initialised with an atom-type embedding, which serves as the initial node feature for subsequent message passing.

**Figure 1:**
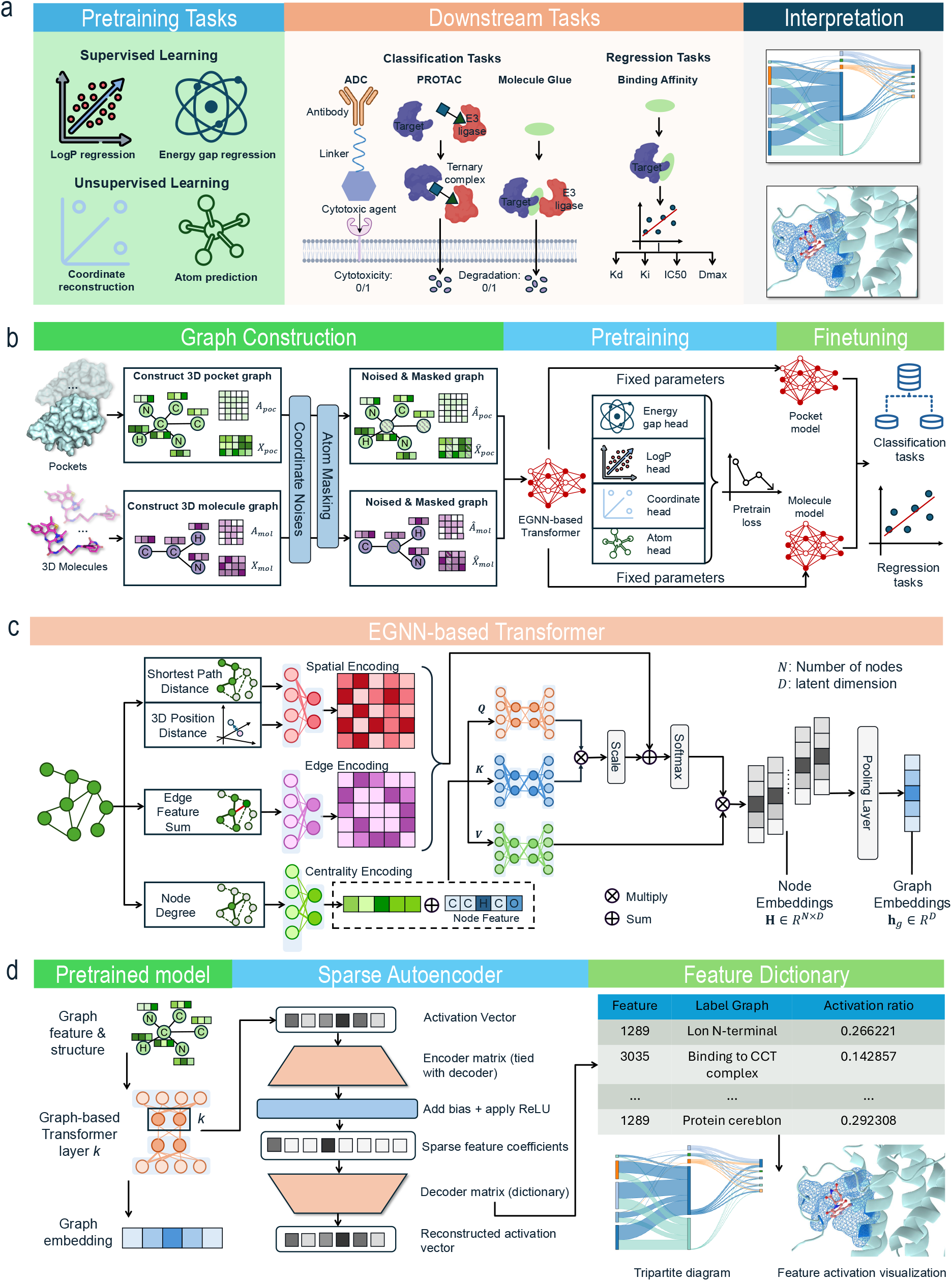
Overview of the MolX framework. **a**. Pretraining objectives, downstream tasks, and interpretation pipeline. MolX is pretrained using supervised objectives (LogP and HOMO–LUMO gap regression) and unsupervised objectives (coordinate reconstruction and atom-type prediction), and subsequently fine-tuned for diverse classification and regression tasks. **b**. Graph construction, pretraining, and fine-tuning workflow for protein pockets and ligand molecules represented as 3D graphs. The number of input ligands and protein pockets can be variable. **c**. E(3)-equivariant graph Transformer architecture integrating centrality, edge, and spatial encodings for geometry-aware message passing and graph-level representation learning. **d**. Sparse autoencoder–based interpretation module that decomposes pretrained model activations into sparse, interpretable features and constructs a feature dictionary linking latent activations to molecular patterns.

MolX captures both structural and chemical attributive information of pocket and ligand graphs through a stack of E(3)-equivariant graph Transformer layers. Within each layer, pocket and ligand representations are updated through a three-stage procedure (Fig. 1c). First, MolX integrates spatial encoding from Graphormer [19] to capture pairwise geometric relationships and edge encoding to represent chemical bond types. Additionally, centrality encoding is adapted to capture node importance and is jointly incorporated with input node features. Second, mutual attention is applied to compute attention coefficients that modulate message passing between atoms. Third, attentional aggregation updates atom-level embeddings, which are subsequently fused with spatial and edge information to produce the final node representations for that layer. Graph-level representations of both pockets and ligands are then obtained by pooling the updated node embeddings.

To enable comprehensive pretraining, MolX is optimised using a combination of unsupervised and supervised learning objectives (Fig. 1b). In the unsupervised learning branch, a coordinate noise masking strategy is applied by randomly perturbing atomic 3D coordinates, and an atom masking strategy is employed by randomly masking atom types in both pocket and ligand graphs. The resulting encoded representations are passed to coordinate reconstruction and atom-type prediction heads to recover the original atomic positions and identities. In the supervised learning branch, two independent multilayer perceptron (MLP) heads are used to predict ligand-level physicochemical properties, specifically the HOMO–LUMO energy gap [37] and LogP [38], from the encoded representations.

After pretraining, the model parameters of MolX are frozen, and the model is fine-tuned for specific downstream tasks. To further enhance model interpretability, MolX incorporates a sparse autoencoder-based interpretation module to analyse internal representations (Fig. 1d). Specifically, we apply a sparse autoencoder to the activation vectors extracted from an intermediate graph Transformer layer. The encoder projects dense activation vectors into a sparse set of feature coefficients by applying a linear transformation followed by a bias term and a ReLU nonlinearity, thereby enforcing sparsity in the latent space. A tied decoder then reconstructs the original activations using a learned dictionary of feature directions. Each learned sparse feature thus corresponds to a distinct and interpretable activation pattern within the pretrained model. By associating these individual sparse features with specific ligand–pocket interaction graphs and quantifying their activation consistency, we construct a feature dictionary that links abstract neural representations to concrete structural motifs. This sparse autoencoder–based analysis provides mechanistic insight into how MolX encodes chemically and structurally meaningful patterns.

### State-of-the-art performance of MolX across diverse pocket-ligand classification benchmarks

We first evaluated the classification performance of MolX on four representative pocket benchmarks, including antibody-drug conjugates (ADC), proteolysis targeting chimeras (PROTAC) [39], molecular glue (MG) [40], and LIT-PCBA, using both scaffold-based [41] and UMAP-based [41] data splits. The UMAP split results are reported in sections S4 and S5 of the Supplementary, and here we focus on the scaffold split results (Tab.1). Across most datasets and evaluation metrics, MolX achieved the best overall performance among all compared methods, including recently proposed foundation models for small-molecule predictions (*e.g.*, MolE, FradNMI, etc.). On the ADC benchmark, MolX reached an AUC of 0.8991, outperforming the strongest baseline AEV-PLIG (0.8939, +0.5 percentage points (pp)). Similar trends were observed for F1 score, where MolX achieved 0.8355, representing a 3.9% relative improvement over the strongest baseline TorchMD-Net (0.8042). These results demonstrate MolX ’s strong ability to learn discriminative ligand–pocket interaction features under scaffold-based generalisation settings.

**Table 1:**
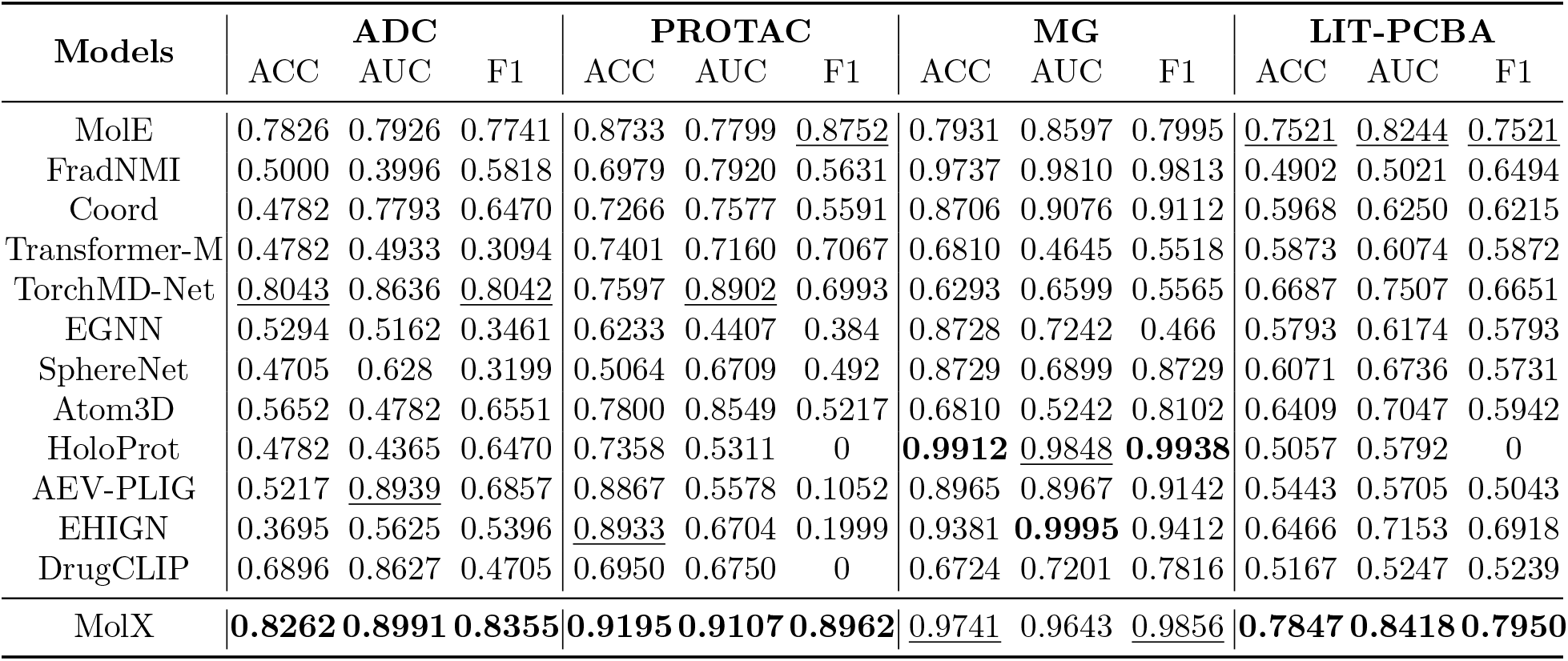
Classification tasks under scaffold split. The best results on each dataset are in bold, the second-to-the-best results are underlined.

| Models | ADC |  |  | PROTAC |  |  | MG |  |  | LIT-PCBA |  |  |
| --- | --- | --- | --- | --- | --- | --- | --- | --- | --- | --- | --- | --- |
|  | ACC | AUC | F1 | ACC | AUC | F1 | ACC | AUC | F1 | ACC | AUC | F1 |
| MolE | 0.7826 | 0.7926 | 0.7741 | 0.8733 | 0.7799 | <u>0.8752</u> | 0.7931 | 0.8597 | 0.7995 | <u>0.7521</u> | <u>0.8244</u> | <u>0.7521</u> |
| FradNMI | 0.5000 | 0.3996 | 0.5818 | 0.6979 | 0.7920 | 0.5631 | 0.9737 | 0.9810 | 0.9813 | 0.4902 | 0.5021 | 0.6494 |
| Coord | 0.4782 | 0.7793 | 0.6470 | 0.7266 | 0.7577 | 0.5591 | 0.8706 | 0.9076 | 0.9112 | 0.5968 | 0.6250 | 0.6215 |
| Transformer-M | 0.4782 | 0.4933 | 0.3094 | 0.7401 | 0.7160 | 0.7067 | 0.6810 | 0.4645 | 0.5518 | 0.5873 | 0.6074 | 0.5872 |
| TorchMD-Net | <u>0.8043</u> | 0.8636 | <u>0.8042</u> | 0.7597 | <u>0.8902</u> | 0.6993 | 0.6293 | 0.6599 | 0.5565 | 0.6687 | 0.7507 | 0.6651 |
| EGNN | 0.5294 | 0.5162 | 0.3461 | 0.6233 | 0.4407 | 0.384 | 0.8728 | 0.7242 | 0.466 | 0.5793 | 0.6174 | 0.5793 |
| SphereNet | 0.4705 | 0.628 | 0.3199 | 0.5064 | 0.6709 | 0.492 | 0.8729 | 0.6899 | 0.8729 | 0.6071 | 0.6736 | 0.5731 |
| Atom3D | 0.5652 | 0.4782 | 0.6551 | 0.7800 | 0.8549 | 0.5217 | 0.6810 | 0.5242 | 0.8102 | 0.6409 | 0.7047 | 0.5942 |
| HoloProt | 0.4782 | 0.4365 | 0.6470 | 0.7358 | 0.5311 | 0 | **0.9912** | <u>0.9848</u> | **0.9938** | 0.5057 | 0.5792 | 0 |
| AEV-PLIG | 0.5217 | <u>0.8939</u> | 0.6857 | 0.8867 | 0.5578 | 0.1052 | 0.8965 | 0.8967 | 0.9142 | 0.5443 | 0.5705 | 0.5043 |
| EHIGN | 0.3695 | 0.5625 | 0.5396 | <u>0.8933</u> | 0.6704 | 0.1999 | 0.9381 | **0.9995** | 0.9412 | 0.6466 | 0.7153 | 0.6918 |
| DrugCLIP | 0.6896 | 0.8627 | 0.4705 | 0.6950 | 0.6750 | 0 | 0.6724 | 0.7201 | 0.7816 | 0.5167 | 0.5247 | 0.5239 |
| MolX | **0.8262** | **0.8991** | **0.8355** | **0.9195** | **0.9107** | **0.8962** | <u>0.9741</u> | 0.9643 | <u>0.9856</u> | **0.7847** | **0.8418** | **0.7950** |

On the more challenging and complex PROTAC benchmark, which involves ternary complex formation and degradation outcomes [39], MolX achieved an AUC of 0.9107, exceeding TorchMD-Net (0.8902, +2.1 pp), the strongest baseline on this metric. The improvement was also observed in F1 score, where MolX reached 0.8962, compared with 0.8752 for MolE and 0.6993 for TorchMD-Net, corresponding to relative improvements of 2.4% and 28.2%, respectively. On the molecular glue benchmark, which reflects small-molecule-induced protein-recruitment and degradation mechanisms exemplified by immunomodulatory drugs [42, 43], MolX remained highly competitive, achieving an AUC of 0.9643 and F1 of 0.9856, with the second-best ACC and F1 among all methods, although HoloProt and EHIGN achieved the best results on selected MG metrics. Notably, MolX substantially outperformed MolE on MG, improving AUC by 10.5 pp and F1 by 18.6 pp. On LIT-PCBA, MolX achieved the best performance across all three metrics, with an AUC of 0.8418 and F1 of 0.7950, outperforming MolE by 1.7 pp in AUC and 4.3 pp in F1. These results highlight MolX ’s advantage in modelling higher-order interaction mechanisms that are central to degradation-based therapeutics [39, 42, 43].

LIT-PCBA is an unbiased dataset for machine learning and virtual screening, providing a stringent benchmark for evaluating whether a model can prioritise active compounds among large candidate pools. To further assess early-recognition capability under the scaffold split, we compared enrichment factors (EFs) at the top 1%, 5%, and 10% of ranked compounds (Tab. 2). MolX achieved the best performance across all three thresholds, with EF@0.01, EF@0.05, and EF@0.10 values of 2.2257, 2.0110, and 1.9485, respectively. Compared with the strongest baseline MolE, MolX improved EF@0.01 from 2.0233 to 2.2257, corresponding to a 10.0% relative gain, while also maintaining consistent advantages at EF@0.05 (+2.5%) and EF@0.10 (+2.5%). Notably, the largest improvement was observed at EF@0.01, indicating that MolX is particularly effective at enriching active compounds within the highest-ranked candidates, a practically important regime for virtual screening where only a small fraction of compounds can be experimentally validated.

**Table 2:** Enrichment factor (EF) results on LIT-PCBA under scaffold split. The best results on each dataset are in bold, the second-to-the-best results are underlined.

| Model | EF@0.01 | EF@0.05 | EF@0.10 |
| --- | --- | --- | --- |
| MolE | <u>2.0233</u> | <u>1.9626</u> | <u>1.9019</u> |
| FradNMI | 0.1065 | 0.0204 | 0.0101 |
| Coord | 1.3843 | 1.3079 | 1.3115 |
| Transformer-M | 1.2139 | 1.4972 | 1.7602 |
| TorchMD-Net | 1.6882 | 1.7143 | 1.6945 |
| EGNN | 1.4910 | 1.4311 | 1.3830 |
| SphereNet | 1.6881 | 1.5972 | 1.5290 |
| Atom3D | 1.9221 | 1.8007 | 1.6995 |
| HoloProt | 1.5174 | 1.5579 | 1.4061 |
| AEV-PLIG | 1.0137 | 1.0948 | 1.0441 |
| EHIGN | 1.8110 | 1.6541 | 1.6192 |
| DrugCLIP | 0.7454 | 0.9809 | 1.0472 |
| MolX | **2.2257** | **2.0110** | **1.9485** |

To evaluate robustness beyond aggregate metrics, we performed subset-level AUC comparisons across PROTAC, MG, and ADC datasets (Fig. 2a). Subsets are defined by target–E3 pairs for

**Figure 2:**
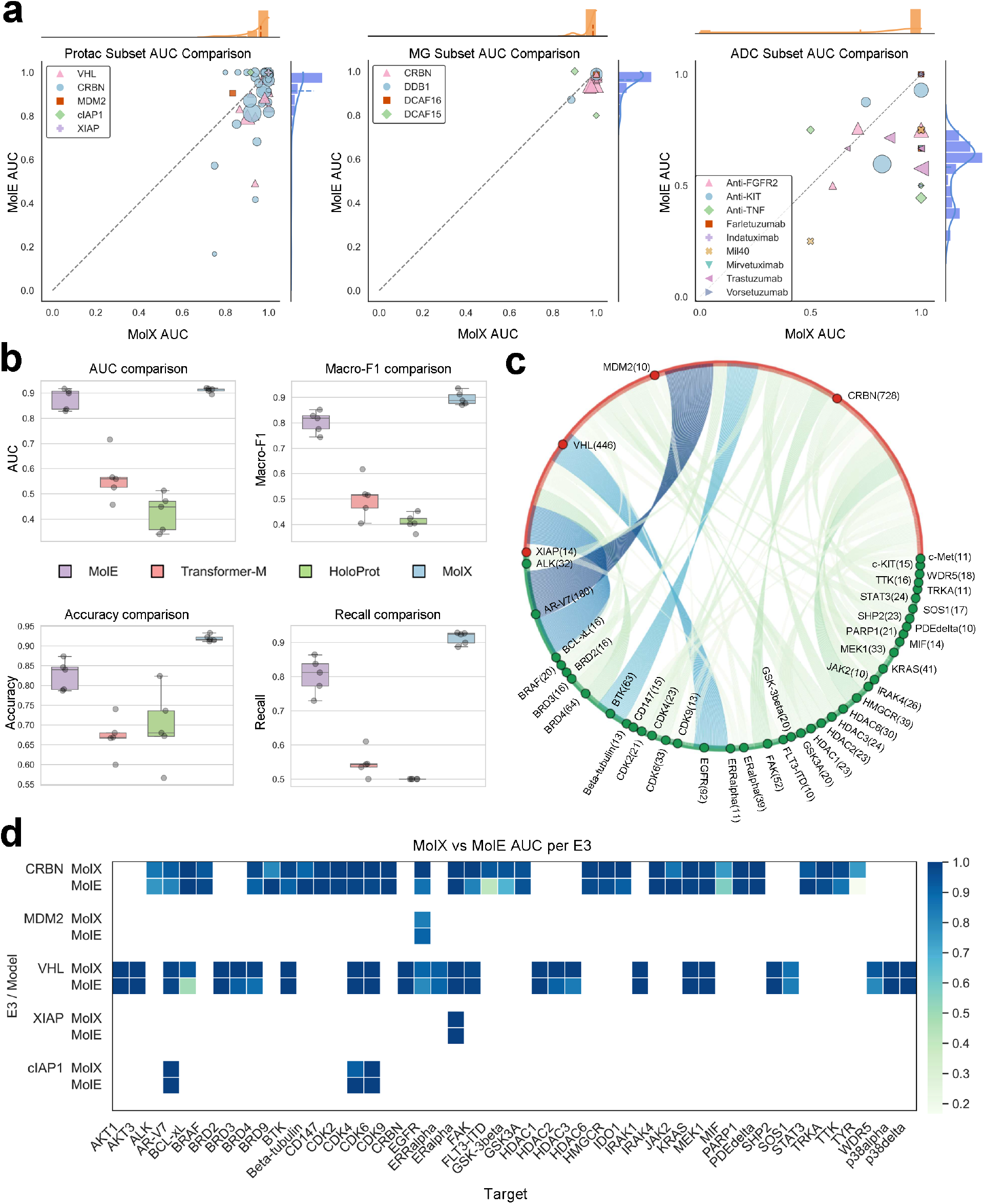
Performance evaluation results on classification tasks under scaffold split. **a**. Subset-level AUC comparison across classification tasks. **b**. Classification performance across multiple metrics. **c**. Fine-grained decomposition of the PROTAC dataset by target–E3 pairs. **d**. Heat map comparison of MolX and MolE across PROTAC target–E3 subsets.

PROTAC and MG, and by antibody–payload pairs for ADC [44]. Pair definitions for PROTAC, MG, and ADC are illustrated in Fig. 2c and Supplementary Figs. S3 and S4, respectively. Each point in Fig. 2a corresponds to a pair-defined subset. The majority of points fall below the diagonal, indicating superior performance of MolX over MolE across the vast majority of subsets. For PROTAC subsets, MolX achieves higher AUCs in over 80% of evaluated E3-specific subsets, with many cases showing absolute gains exceeding 0.15. Similar dominance patterns are observed for MG E3-based subsets and ADC antibody-based subsets, confirming that MolX ’s advantages generalise broad robustness across biological contexts rather than being driven by a small number of favourable cases.

Beyond AUC, MolX also delivers consistent gains across metrics including accuracy, recall, and macro-F1 scores (Fig. 2b). Averaged across classification tasks, MolX improves macro-F1 by approximately 12–18 pp relative to Transformer-M and HoloProt, and by 8–10 pp over MolE. MolX also demonstrates the lowest standard errors in AUC and accuracy among all evaluated models, indicating more stable performance across folds. Notably, MolX achieves the highest recall values, underscoring strong sensitivity to positive degradation events, which is particularly critical for early-stage screening tasks. These improvements suggest that MolX not only ranks positives more effectively but also yields more balanced decision boundaries.

We further analysed MolX’s performance on the PROTAC dataset by stratifying samples into 61 target–E3 pair subsets (Fig. 2c). This fine-grained decomposition reveals substantial heterogeneity across pairs. Importantly, MolX consistently outperforms MolE across the majority of these pairs. As visualised in the AUC heat map (Fig. 2d), MolX shows clear advantages for major E3 ligases such as VHL and CRBN, where average AUC improvements over MolE exceed 0.20 for several targets (*e.g.*, BRD4, CDK6, ER*α*). Even for sparse or challenging pairs such as EGFR-MDM2 or ER*α*-XIAP, MolX maintains competitive performance, indicating superior generalisation under data scarcity.

The suboptimal performance of existing classification models largely stems from their limited ability to capture interaction-level structural context. Most geometry-aware or ligand-centric approaches typically focus on local atomic features or isolated molecular representations, while failing to jointly model protein pockets and ligands to capture higher-order interaction patterns that are critical for degradation-related tasks. MolX addresses these limitations by jointly encoding ligands and protein pockets as 3D graphs within an E(3)-equivariant Transformer framework, enabling geometry-aware, interaction-centric representation learning. As a consequence, MolX learns more transferable and discriminative representations, leading to consistent and substantial performance gains across classification benchmarks and fine-grained biological subsets.

### Consistent generalisation of MolX on regression and property prediction tasks

We also evaluated MolX across a diverse suite of regression tasks, including binding affinity prediction (*K_d_, K_i_, IC*_50_) using the PDBbind dataset [45] and physicochemical property prediction (*EA, EN, η, IP, KPT* , and *MW* ) using the MISATO benchmark [46]. For PDBbind, we evaluated model performance under the scaffold split on the general set and also trained models on the PDBbind-refined set, using CASF-16 [47], OODTest [48], and FEP [49] as external testing benchmarks. Performance was compared against a range of state-of-the-art baselines (Tables 3, 4, and 5). The results under UMAP split [41] are reported in sections S4 and S5 in the Supplementary Information. Across these evaluated regression tasks, MolX consistently achieved the lowest or near-lowest mean absolute error (MAE) and root mean square error (RMSE), while also showing strong rank-correlation performance, demonstrating robust generalisation across both interaction-centric and intrinsic molecular property regression settings.

**Table 3:** Model performance on PDBbind dataset under scaffold split. The best results on each dataset are in bold, the second-to-the-best results are underlined.

| Models | Kd |  |  | Ki |  |  | IC50 |  |  |
| --- | --- | --- | --- | --- | --- | --- | --- | --- | --- |
|  | MAE | RMSE | Spearman | MAE | RMSE | Spearman | MAE | RMSE | Spearman |
| MolE | 1.2467 | <u>1.5066</u> | 0.4675 | 1.4706 | 1.8630 | 0.5877 | <u>1.0657</u> | <u>1.3560</u> | <u>0.6640</u> |
| FradNMI | 1.7508 | 2.4411 | 0.2797 | 1.9094 | 2.4127 | 0.4426 | 1.6028 | 2.0528 | 0.4323 |
| Coord | 1.4906 | 1.9558 | 0.3224 | 1.5561 | 2.0014 | 0.5116 | 1.2469 | 1.5841 | 0.5018 |
| Transformer-M | 1.3641 | 1.7767 | 0.5086 | 1.5622 | 2.0424 | 0.5702 | 1.1586 | 1.4701 | 0.4964 |
| TorchMD-Net | **1.1820** | 1.5144 | <u>0.5361</u> | <u>1.2970</u> | <u>1.6181</u> | <u>0.6167</u> | 1.2319 | 1.5771 | 0.4916 |
| EGNN | 1.4059 | 1.7728 | 0.2913 | 1.5788 | 2.0213 | 0.2895 | 1.336 | 1.6721 | 0.3071 |
| SphereNet | 4.3857 | 4.7218 | 0.2169 | 4.4942 | 4.9033 | 0.2143 | 5.1579 | 5.4288 | 0.2569 |
| Atom3D | 1.4476 | 1.8309 | -0.0420 | 1.7573 | 2.1726 | 0.1393 | 1.4210 | 1.7757 | 0.0935 |
| HoloProt | 1.2260 | 1.5481 | 0.3927 | 1.3572 | 1.7801 | 0.5854 | 1.0896 | 1.3729 | 0.6093 |
| AEV-PLIG | 1.2586 | 1.6251 | 0.2922 | 1.5483 | 1.9284 | 0.5631 | 1.2592 | 1.5629 | 0.5379 |
| EHIGN | 1.2870 | 1.6471 | 0.3639 | 1.5064 | 1.9530 | 0.4500 | 1.1635 | 1.4512 | 0.5656 |
| DrugCLIP | 1.2507 | 1.5696 | 0.3631 | 1.4065 | 1.8018 | 0.5508 | 1.1156 | 1.3778 | 0.5498 |
| MolX | <u>1.2007</u> | **1.4607** | **0.6394** | **1.2810** | **1.6106** | **0.6616** | **1.0319** | **1.3385** | <u>0.6567</u> |

**Table 4:** Model performance on PDBbind-refined dataset with CASF-16, OODTest and FEP as testing sets. The best results on each dataset are in bold, the second-best results are underlined.

| Models | CASF-16 |  |  | OODTest |  |  | FEP |  |  |
| --- | --- | --- | --- | --- | --- | --- | --- | --- | --- |
|  | MAE | RMSE | Spearman | MAE | RMSE | Spearman | MAE | RMSE | Spearman |
| MolE | 1.8505 | 2.1010 | 0.4081 | 1.3931 | 1.6964 | 0.5521 | 13.4231 | 13.5820 | -0.0878 |
| FradNMI | 2.0657 | 2.8711 | 0.5289 | 1.9499 | 2.8052 | 0.4324 | <u>6.0654</u> | <u>7.6622</u> | -0.0291 |
| Coord | 1.3857 | 1.7380 | 0.6218 | 1.2777 | 1.6317 | 0.6345 | 25.5991 | 25.9591 | -0.2739 |
| Transformer-M | <u>1.2143</u> | <u>1.5807</u> | <u>0.6920</u> | 1.2993 | 1.6683 | 0.6152 | 13.5436 | 13.7547 | -0.0917 |
| TorchMD-Net | 1.2878 | 1.6017 | 0.6325 | <u>1.2004</u> | <u>1.5059</u> | 0.6308 | 15.8966 | 16.0847 | **0.4187** |
| EGNN | 1.6407 | 1.9844 | 0.2686 | 1.5429 | 1.8598 | 0.3243 | 9.3078 | 9.4812 | 0.0059 |
| SphereNet | 1.5559 | 1.9023 | 0.3433 | 1.5122 | 1.8404 | 0.3961 | 9.2987 | 9.4723 | 0.1313 |
| Atom3D | 1.4529 | 1.8163 | 0.5473 | 1.4060 | 1.7754 | 0.4395 | 16.3923 | 16.5405 | -0.2978 |
| HoloProt | 1.3392 | 1.7048 | 0.6362 | 1.4725 | 1.9137 | 0.4910 | 14.4644 | 14.6105 | -0.2221 |
| AEV-PLIG | 1.7809 | 2.2228 | 0.2144 | 1.8481 | 2.6493 | 0.4085 | 14.6764 | 15.0939 | -0.1449 |
| EHIGN | 1.8933 | 2.2363 | 0.6793 | 1.6497 | 1.9707 | <u>0.6579</u> | 16.5870 | 16.7569 | -0.4198 |
| DrugCLIP | 1.4329 | 1.7856 | 0.5648 | 1.4992 | 1.8132 | 0.4846 | 16.5702 | 16.7212 | -0.2822 |
| MolX | **1.1697** | **1.4815** | **0.7156** | **0.9639** | **1.2644** | **0.6691** | **5.4688** | **5.6082** | <u>0.3102</u> |

On binding affinity prediction tasks under the PDBbind scaffold split, MolX achieved the best overall performance across most metrics. For *K_d_* prediction, MolX reduced RMSE to 1.4607, outperforming the strongest baseline MolE (1.5066, 3.0% reduction), and achieved the highest Spearman correlation of 0.6394, exceeding TorchMD-Net (0.5361). Although TorchMD-Net obtained the lowest MAE for *K_d_*, MolX remained the second-best method with an MAE of 1.2007. Similar trends were observed for *K_i_*, where MolX achieved the best MAE, RMSE, and Spearman correlation, reducing MAE to 1.2810 and RMSE to 1.6106 compared with the strongest baseline TorchMD-Net (1.2970 MAE and 1.6181 RMSE). Notably, for *IC*_50_ prediction, MolX achieved the lowest MAE and RMSE, with an MAE of 1.0319 and RMSE of 1.3385, corresponding to reductions of 3.2% and 1.3% relative to MolE, respectively, while maintaining the second-best Spearman correlation. These results indicate MolX’s superior ability to capture fine-grained ligand–pocket interaction patterns critical for quantitative affinity estimation.

To further assess out-of-distribution generalisation, we trained models on the PDBbind-refined set and evaluated them on CASF-16, OODTest, and FEP benchmarks (Table 4). MolX achieved the best MAE, RMSE, and Spearman correlation on both CASF-16 and OODTest, with particularly strong gains on OODTest, where it reduced MAE and RMSE to 0.9639 and 1.2644, respectively. On the FEP benchmark, MolX achieved the lowest MAE and RMSE, reducing MAE to 5.4688 and RMSE to 5.6082, outperforming the strongest error-based baseline, FradNMI, by 9.8% and 26.8%, respectively. Although TorchMD-Net achieved the highest Spearman correlation on FEP, MolX ranked second, suggesting that MolX provides strong absolute affinity accuracy while retaining competitive ranking ability in challenging external benchmarks.

MolX also demonstrated strong performance on physicochemical property prediction tasks, achieving the lowest MAE on five of six evaluated properties in the MISATO benchmark and the second-best MAE on molecular weight prediction (Table 5). In particular, MolX reduced MAE to 0.1352 for EA, outperforming the strongest baseline DrugCLIP (0.1584, 14.6% reduction), and achieved the best RMSE of 0.2739. Similar gains were observed for EN, *η*, IP, and KPT, where MolX achieved MAE values of 0.1832, 0.0435, and 0.0842, respectively, and obtained the best RMSE for *η* and KPT. While geometry-aware models such as TorchMD-Net, EGNN, and SphereNet perform competitively on selected physicochemical properties, their performance is less consistent on binding affinity tasks that require explicit ligand-pocket interaction modelling. Conversely, large-scale pretrained approaches such as FradNMI show limited regression accuracy across both affinity and property prediction settings. MolX bridges this gap by jointly encoding ligand and pocket graphs within a unified E(3)-equivariant framework and by leveraging hybrid supervised + self-supervised pretraining objectives, enabling it to outperform both geometry-focused and representation-centric baselines across different regression settings.

**Table 5:**
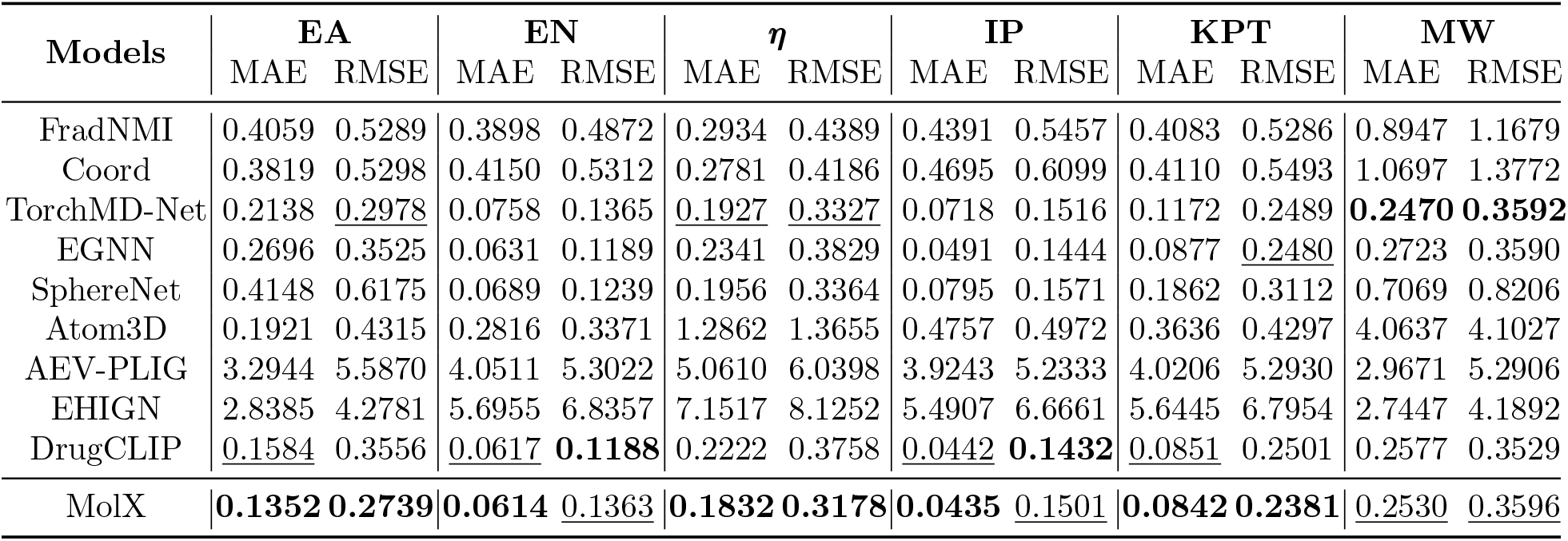
Model performance on MISATO dataset under scaffold split, EA, EN, *η*, IP, KPT and MW is the abbreviation for Electron Affinity, Electronegativity, Hardness, Ionisation Potential, Koopman value, and Molecular Weight, respectively.

| Models | EA | | EN | | $\eta$ | | IP | | KPT | | MW | |
| --- | --- | --- | --- | --- | --- | --- | --- | --- | --- | --- | --- | --- |
|  | MAE | RMSE | MAE | RMSE | MAE | RMSE | MAE | RMSE | MAE | RMSE | MAE | RMSE |
| FradNMI | 0.4059 | 0.5289 | 0.3898 | 0.4872 | 0.2934 | 0.4389 | 0.4391 | 0.5457 | 0.4083 | 0.5286 | 0.8947 | 1.1679 |
| Coord | 0.3819 | 0.5298 | 0.4150 | 0.5312 | 0.2781 | 0.4186 | 0.4695 | 0.6099 | 0.4110 | 0.5493 | 1.0697 | 1.3772 |
| TorchMD-Net | 0.2138 | <u>0.2978</u> | 0.0758 | 0.1365 | <u>0.1927</u> | <u>0.3327</u> | 0.0718 | 0.1516 | 0.1172 | 0.2489 | **0.2470** | **0.3592** |
| EGNN | 0.2696 | 0.3525 | 0.0631 | 0.1189 | 0.2341 | 0.3829 | 0.0491 | 0.1444 | 0.0877 | <u>0.2480</u> | 0.2723 | 0.3590 |
| SphereNet | 0.4148 | 0.6175 | 0.0689 | 0.1239 | 0.1956 | 0.3364 | 0.0795 | 0.1571 | 0.1862 | 0.3112 | 0.7069 | 0.8206 |
| Atom3D | 0.1921 | 0.4315 | 0.2816 | 0.3371 | 1.2862 | 1.3655 | 0.4757 | 0.4972 | 0.3636 | 0.4297 | 4.0637 | 4.1027 |
| AEV-PLIG | 3.2944 | 5.5870 | 4.0511 | 5.3022 | 5.0610 | 6.0398 | 3.9243 | 5.2333 | 4.0206 | 5.2930 | 2.9671 | 5.2906 |
| EHIGN | 2.8385 | 4.2781 | 5.6955 | 6.8357 | 7.1517 | 8.1252 | 5.4907 | 6.6661 | 5.6445 | 6.7954 | 2.7447 | 4.1892 |
| DrugCLIP | <u>0.1584</u> | 0.3556 | <u>0.0617</u> | **0.1188** | 0.2222 | 0.3758 | <u>0.0442</u> | **0.1432** | <u>0.0851</u> | 0.2501 | 0.2577 | 0.3529 |
| MolX | **0.1352** | **0.2739** | **0.0614** | <u>0.1363</u> | **0.1832** | **0.3178** | **0.0435** | <u>0.1501</u> | **0.0842** | **0.2381** | <u>0.2530</u> | <u>0.3596</u> |

Across binding affinity and physicochemical property regression tasks, MolX generally achieves the lowest errors and highest rank correlations compared with all baselines (Fig. 3a). As shown in Fig. 3a, MolX reduces both MAE and RMSE while improving Spearman correlation for representative properties such as electron affinity in MISATO, and binding affinities in CASF-16 and OODTest, indicating more accurate and monotonic predictions. The radial comparison in Fig. 3b further demonstrates MolX’s robustness across diverse datasets, including multiple PDBbind-derived affinity benchmarks, where competing models exhibit pronounced variability in performance. Fig. 3c shows that MolX yields a tighter alignment with the identity line and reduced noise compared with FradNMI and Atom3D. To assess stability across the target range, Fig. 3d groups samples into equal-width bins based on ground-truth values and compares average prediction trends within each bin. MolX maintains stable and well-calibrated predictions across bins, particularly at distribution extremes where baseline models degrade. A comprehensive summary across all tasks and metrics (Fig. 3e) consolidates that MolX consistently outperforms geometry-based and foundation-model baselines, highlighting its effectiveness as a unified regression framework for both interaction-driven and intrinsic molecular property prediction.

**Figure 3:**
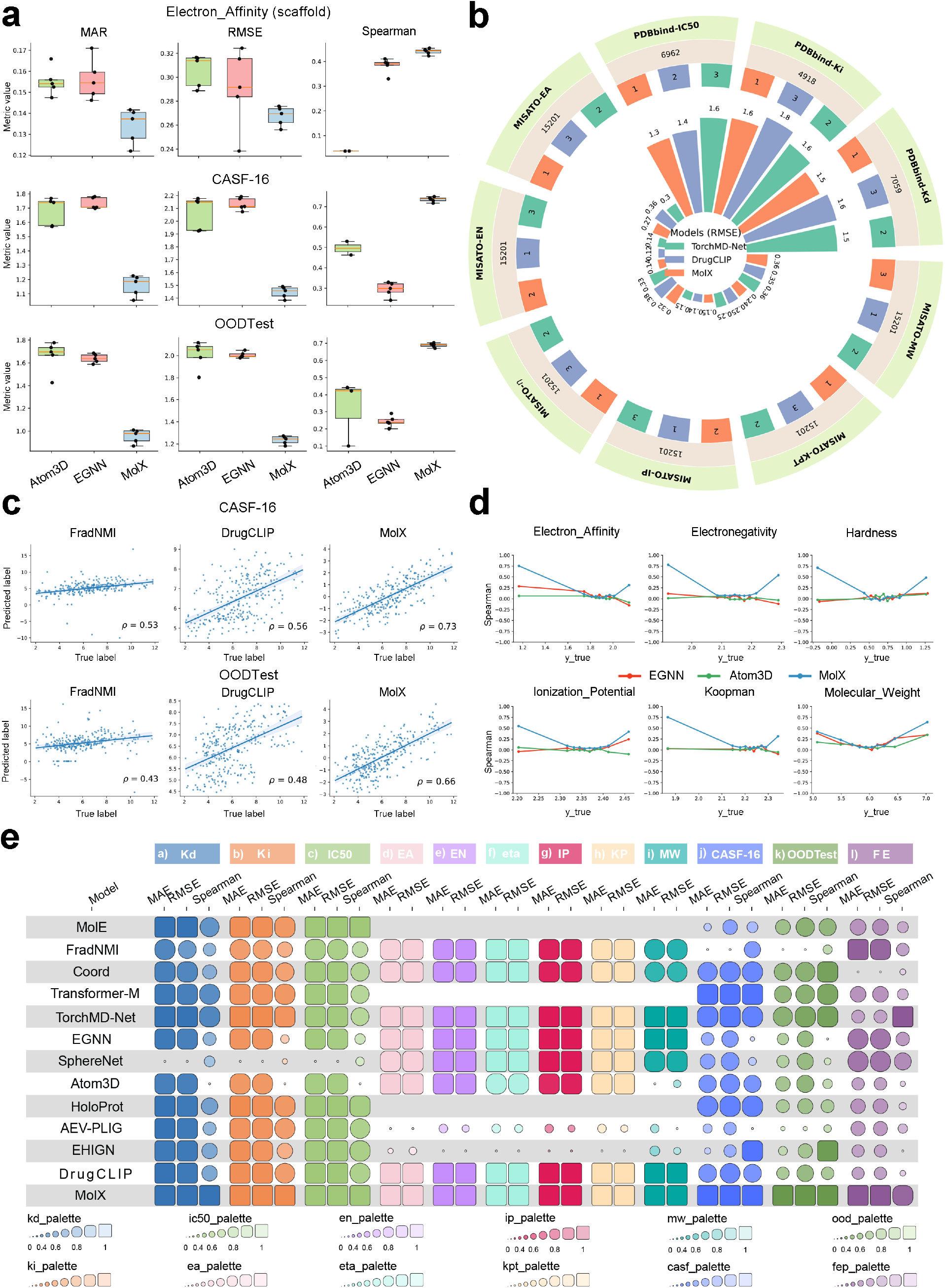
Regression results under the scaffold split. **a**. Comparison of MAE, RMSE, and Spearman correlation for predictions of electron affinity and binding affinities across representative models. **b**. Radial bar plot comparing RMSE across multiple regression tasks, including PDBbind and MISATO datasets. **c**. Scatter plots of predicted versus ground-truth values for regression of pdbbind CASF-16 and OODTest, comparing FradNMI, Atom3D, and MolX. **d**. Sensitivity analysis showing prediction trends across binned target ranges for multiple regression properties. **e**. Summary heatmap comparing regression performance (MAE, RMSE, and Spearman) of all models across pdbbind and MISATO. Larger rectangles indicate better relative performance for each metric and task.

### Spatial bias of geometry-enhanced model capability

Unlike recently proposed molecular foundation models [6, 17, 18] which employ Transformers [23] as the core backbone to encode molecule or pocket structures, the key innovation of MolX lies in introducing a spatial position bias to the original Transformer that explicitly encodes 3D geometric dependencies between atoms. As discussed in [19], although standard Transformers have a global receptive field in representation learning, they need to explicitly specify different positions or encode the positional dependency. In molecular systems, however, atoms are not ordered sequentially but instead occupy coordinates in continuous three-dimensional space, where geometric proximity rather than sequence adjacency governs interactions. To address this, MolX integrates a spatial positional bias into the attention mechanism, allowing the attention weights to be modulated by pairwise Euclidean distances between atoms, thereby coupling attention strength with real spatial geometry.

To illustrate this behaviour, Fig. 4 visualises representative ligands exhibiting distinct spatial configurations. Atoms that are located in spatially distant regions of the molecular graph due to ring substituents (Fig. 4a), elongated conformation (Fig. 4b) and rotatable bonds (Fig. 4c) exhibit larger Euclidean distances, as highlighted by the red regions in the leftmost 3D molecule structures and the red frames in the corresponding distance heatmaps. In these cases, the learned spatial bias term yields proportionally reduced attention scores for distant atoms, enabling the model to prioritise geometrically relevant local interactions while still being able to capture global structural determinants. This bias, therefore, acts as a geometry-aware gating mechanism, ensuring that information propagation reflects real 3D structure rather than topological connectivity only.

**Figure 4:**
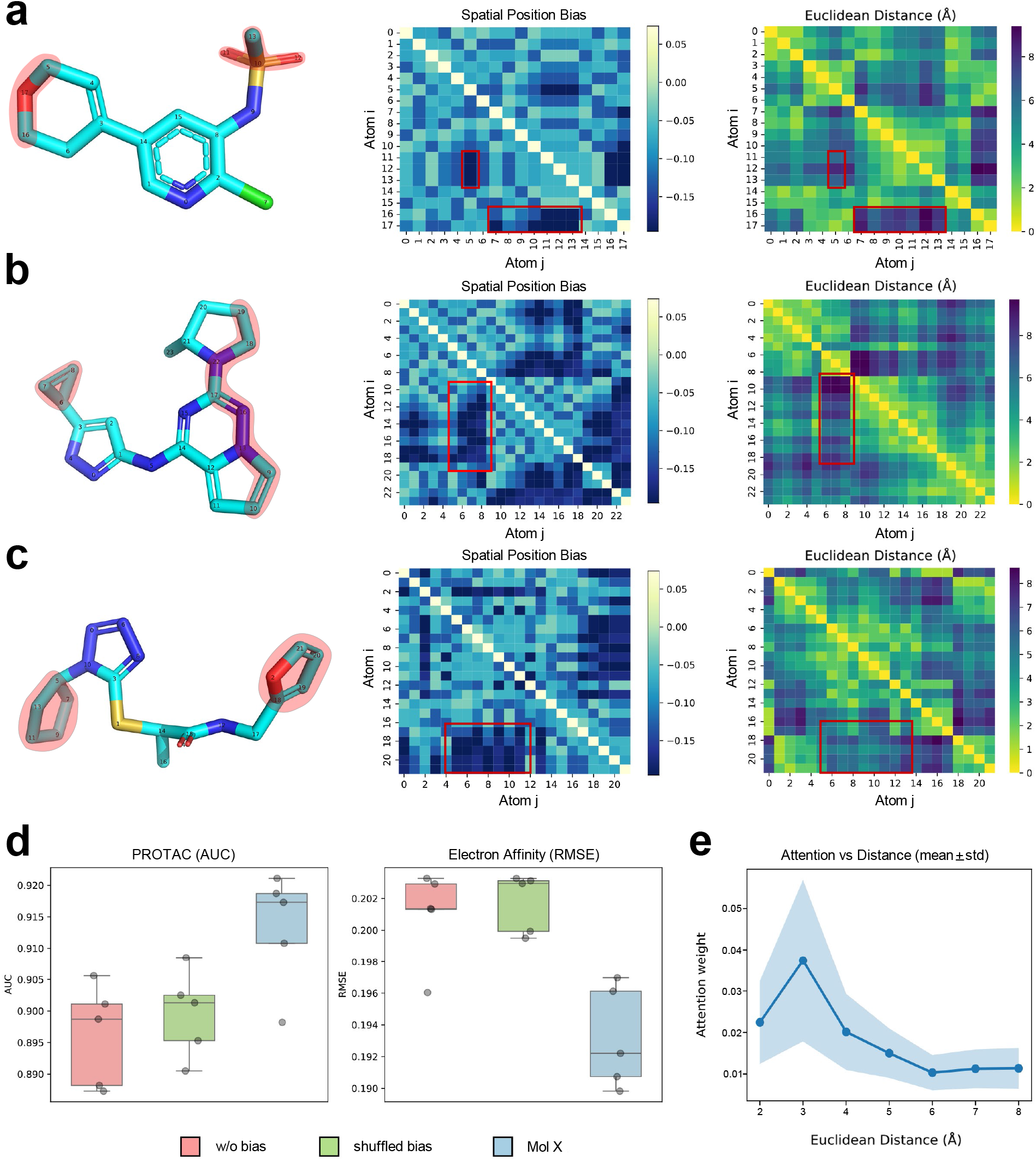
The analysis of 3D positional spatial encoding. The three ligands are representative examples selected from the PDBbind dataset to illustrate distinct and commonly encountered molecular geometries in protein-ligand complexes. **a**. The ligand contains atom pairs that are spatially distant due to ring substituents extending away from the molecular core. **b** The ligand adopts a more elongated conformation, leading to pronounced long-range spatial separations between atoms. **c**. The ligand is characterised by increased conformational flexibility, with multiple rotatable bonds giving rise to non-planar geometries. **d**. Performance comparison under spatial-bias ablation. **e**. The trend of attention weight versus inter-atomic distance on the PROTAC dataset.

To quantify the contribution of spatial bias, we performed an ablation study by removing (w/o bias) or randomly shuffling (shuffled bias) the bias term in the E(3)-equivariant Transformer layers. Across both classification and regression benchmarks, both ablated variants exhibit a consistent decline in performance compared to the full MolX model (Fig. 4d). Furthermore, Fig. 4e reveals that the biased model concentrates substantially higher attention on geometrically proximal atom pairs (≤ 3Å), with attention weights peaking at short inter-atomic distances and decaying smoothly as distance increases.

From a biochemical perspective, by inducing a smooth distance-dependent attenuation of attention instead of imposing a hard cutoff, MolX remains sensitive to long-range geometric dependencies that are crucial for describing flexible or non-planar ligands, where non-local atomic interactions can dominate functional outcomes. By aligning the attention mechanism with physical geometry, MolX achieves a closer correspondence between model saliency and experimentally known binding determinants, enhancing both predictive accuracy and interpretability.

### 3D Coordinate noising strengthens geometry-sensitive pretraining

Another key innovation of MolX lies in its 3D coordinate noising module, which plays a crucial role in enabling geometry-sensitive self-supervised learning. To investigate its contribution in the pretraining process, we performed a comprehensive ablation and qualitative analysis, as summarised in Fig. 5. Figure 5a presents ablation study results across multiple classification and regression benchmarks, where individual pretraining objectives are selectively removed. In particular, w/o-3d denotes the removal of the 3D coordinate masking and reconstruction loss, while w/o-atom corresponds to removing the atom-type prediction objective. Performance is reported using AUC for classification tasks (Protac, MG, ADC, PCBA) and RMSE for regression tasks (PDBbind-Kd and MISATO properties). Removing the 3D coordinate noising objective consistently results in the most pronounced performance degradation across almost all tasks, larger than the drops observed when excluding atom-type supervision or ligand physicochemical objectives. This trend indicates that learning to recover corrupted atomic coordinates provides a dominant self-supervised signal, particularly for downstream tasks that rely on precise geometric reasoning.

**Figure 5:**
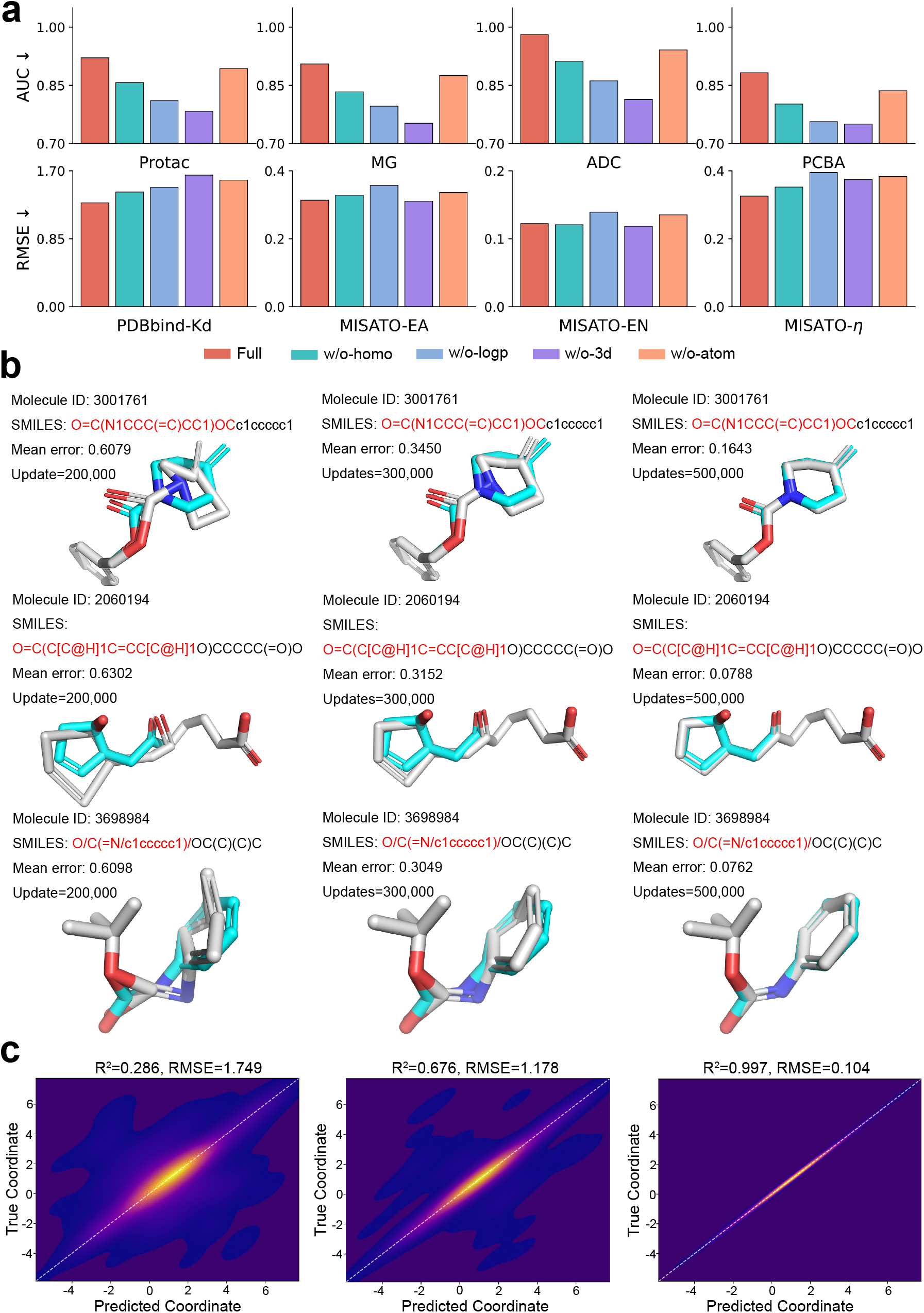
Loss funtion analysis. **a**. Ablation results on classification (AUC) and regression (RMSE) tasks under different pretraining objectives. **b**. Representative molecules with predicted atomic coordinates at increasing pretraining steps. **c**. Density plots of predicted versus true atomic coordinates at different training stages.

Figure 5b visualises the coordinate noising and denoising process during pretraining for three representative small molecules. Atomic coordinates are first perturbed with Gaussian noise at the input and then progressively reconstructed by the model as training proceeds. The visualisations show molecular conformations at different training checkpoints (200k, 300k, and 500k updates), where the predicted atomic positions gradually converge steadily toward the ground-truth geometries. Mean coordinate errors decrease monotonically with training, demonstrating that the model effectively learns to internalise physically consistent geometric priors rather than merely memorising local atom neighbourhoods.

To quantitatively assess the reconstruction quality, Figure 5c plots predicted versus true atomic coordinates across three representative stages or model variants, visualised using density-based heatmaps. Each point corresponds to one atomic coordinate dimension, and the dashed diagonal denotes perfect reconstruction. Without effective coordinate noise, predictions exhibit substantial dispersion and weak correlation (R² = 0.286, RMSE = 1.749). As training proceeds and the coordinate reconstruction objective is fully exploited, predictions align tightly with the identity line, achieving a near-perfect fit (R² = 0.997, RMSE = 0.104). These results confirm that 3D coordinate noising enforces a strong geometric consistency constraint, enabling the model to learn smooth, globally coherent spatial representations that respect molecular geometry.

Taken together, the ablation results, qualitative trajectory visualisations, and quantitative coordinate regression analyses collectively demonstrate that 3D coordinate noising is a key driver of MolX’s self-supervised learning effectiveness. By explicitly training the model to denoise corrupted atomic geometries, this objective introduces a physically meaningful inductive bias into the pretrained representations, which in turn yields consistent performance gains across both classification and regression tasks. This mechanism ensures that the pretrained model captures not just topological connectivity but the continuous spatial manifold underlying molecular and pocket structures, thereby improving transferability to geometry-intensive downstream tasks.

### Geometry-aware interpretability of MolX at the pocket level

To elucidate the internal representations learned by MolX, we leverage the sparse autoencoder (SAE) module to analyse the dictionary-layer activation features that underlie protein-ligand interaction predictions (Fig. 6a). The middle column in Fig. 6a displays a subset of sparse activation features sampled from a 4096-dimensional dictionary, where each feature represents a distinct and interpretable activation pattern extracted from a pretrained MolX Transformer layer. In contrast to dense embeddings, these sparse features are selectively activated, enabling a direct bridge between model latent neural representations and biologically meaningful interaction descriptors.

**Figure 6:**
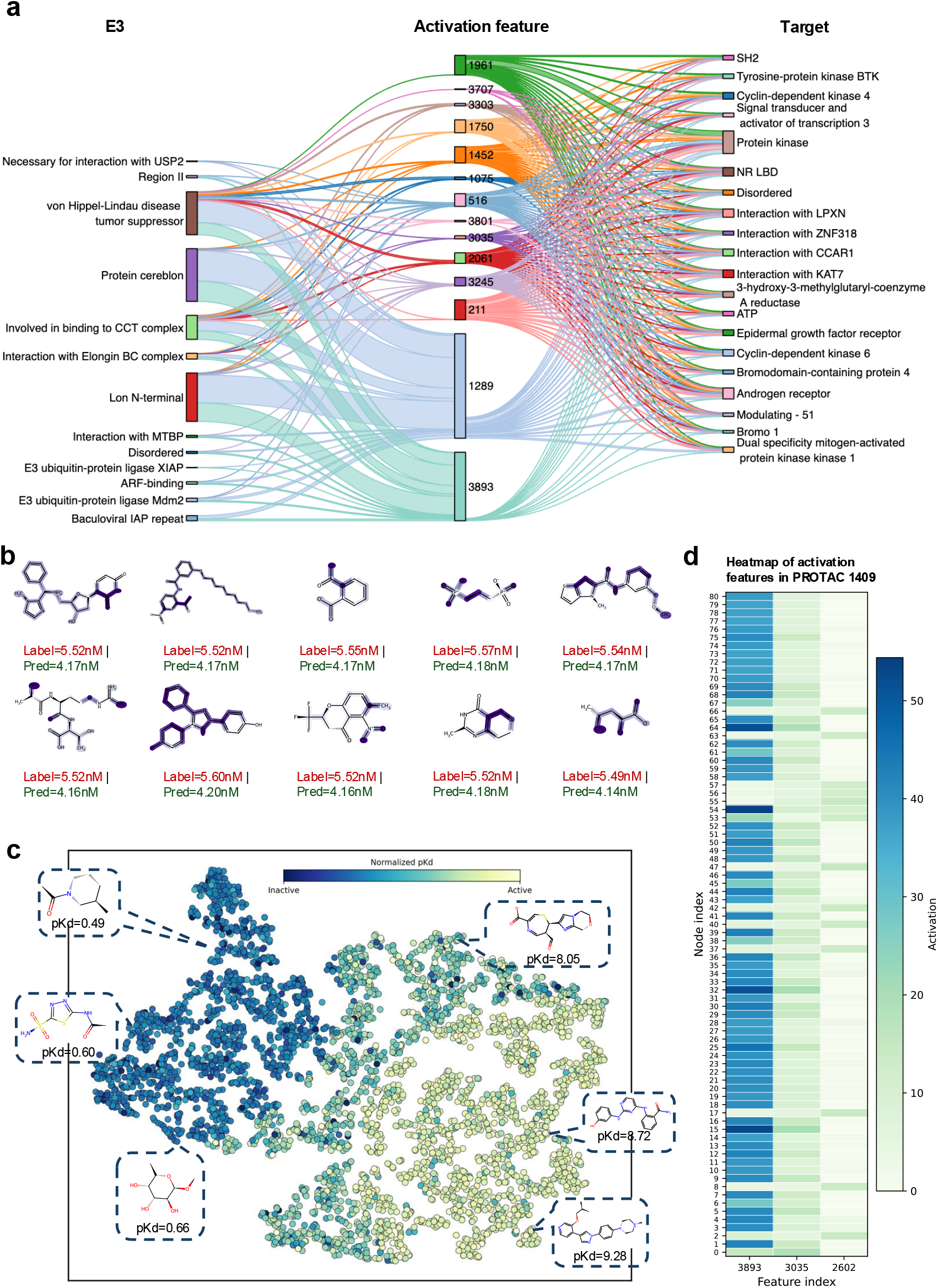
Interpretability analysis of MolX using sparse autoencoder features. **a**. Tripartite diagram linking E3 descriptors, shared activation features, and target descriptors, revealing interpretable interaction motifs captured by sparse autoencoder dictionary features. **b**. Atom-level activation visualisation for representative small molecules from the PDBbind test set, where atoms are colored by their mean sparse autoencoder activation values, highlighting chemically relevant substructures. **c**. t-SNE projection of activation features for PDBbind test-set molecules, with points colored by normalised *pKd* values (darker colours indicate higher activity). **d**. Heatmap of activation values for PROTAC 1409 across the three highest-mean activation feature dimensions.

The left and right columns correspond to E3 ligase–specific and target protein–specific descriptors, respectively. These descriptors are derived directly from UniProt [50] annotations by parsing protein text records (e.g., DOMAIN, REGION, CHAIN, and BINDING fields). Each descriptor is mapped to a specific amino-acid range, such as a functional domain or binding region [51, 52]. For a given protein pocket, residues falling within the annotated range are identified, and the mean activation values of each SAE feature over these residues are computed. This procedure establishes a quantitative link between internal sparse activation features and known protein structural or functional annotations.

To highlight the most informative associations, activation features are ranked for each descriptor according to their mean activation values, and only the top 10% of features are retained. Furthermore, we restrict the analysis to activation features that are shared between both the E3 and target sides, ensuring that the resulting features capture interaction-relevant patterns rather than protein-specific artefacts. The resulting tripartite network forms the basis of the visualisation in Fig. 6a. From this analysis, several biologically meaningful patterns emerge. Notably, multiple activation features simultaneously connect E3 ligase functional regions, such as the von Hippel–Lindau (VHL) interaction interface [53], cereblon-associated regions [54], and Elongin BC complex–binding motifs together with target protein domains including kinase domains [55], bromodomains [56], nuclear receptor ligand-binding domains [57], as well as transcriptional regulators (*e.g.*, STAT3 [58] and androgen receptor [59]). These co-activation patterns suggest that MolX learns high-level abstract feature representations that encode interaction compatibility between E3 machinery and target functional modules, a critical property for modelling PROTAC-mediated degradation.

In addition, several activation features correspond to disordered regions and experimentally known protein–protein interaction interfaces, particularly on the E3 side. This observation aligns with experimental evidence that intrinsically disordered or flexible regions often facilitate ternary complex formation and ubiquitination efficiency [60, 61]. The fact that such regions emerge naturally from SAE analysis, without being explicitly labelled during training, highlights MolX’s capacity to internalise higher-order structural and functional priors relevant to PROTAC activity.

Importantly, the sparse autoencoder not only highlights correlations but also provides a mechanistic decomposition of MolX’s internal decision process. Each dictionary feature corresponds to a reusable, localised activation pattern that can be traced back to specific protein regions on both the E3 and target sides. This enables interpretation at the level of which interaction motifs are being jointly recognised, rather than merely highlighting which residues are important. This mechanistic interpretability is further supported by a concrete structural example shown in Fig. 7a, which visualises the spatial relationship between the target protein BCL2 [62, 63] and a representative PROTAC (compound 272). In this figure, the binding pocket of BCL2 is highlighted and coloured based on the mean activation values of the corresponding sparse autoencoder dictionary features, aggregated at the residue level. Residues with higher activation are rendered in deeper blue, indicating stronger engagement of the learned activation patterns. Notably, the regions exhibiting the highest activation concentrate around the ligand-binding pocket that directly accommodates the PROTAC, rather than being uniformly distributed across the protein surface. This spatial localisation demonstrates that the SAE-derived features are not abstract or diffuse signals, but instead correspond to geometrically coherent interaction motifs that align with known binding interfaces. The close correspondence between high-activation residues and the PROTAC-contacting pocket provides direct structural evidence that MolX internal representations encode functionally relevant interaction patterns.

**Figure 7:**
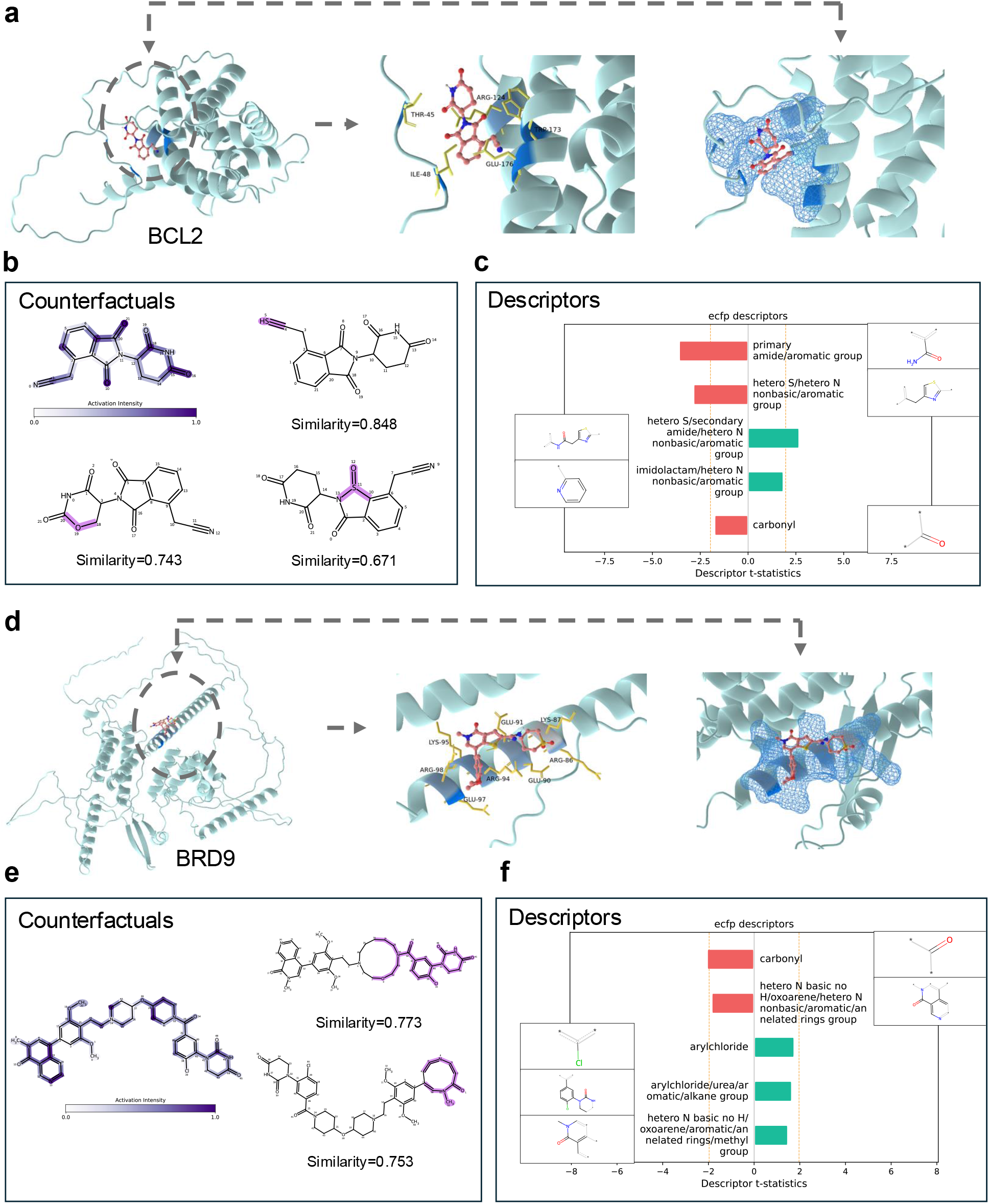
Case studies on PROTACs targeting BCL2 [62] and BRD9 [64]. **a**. visualisations of the spatial relationship between the PROTAC and the BCL2 binding pocket, where pocket residues are colored by their mean sparse autoencoder activation values (blue intensity indicates higher activation). **b**. The counterfactual PROTAC molecules generated by substituting functional groups at highly activated atomic regions, with structural similarity to the original compound indicated. **c**. Summaries of descriptor-level contributions inferred from counterfactual perturbations, highlighting functional groups that positively or negatively influence model predictions. **d**. Case study for target protein BRD9. **e** Counterfactual PROTACs targeting BRD9. **f**. Descriptors inferred for the PROTAC targeting BRD9.

### Chemical interpretability of MolX at the molecular level

Beyond protein-centric interpretation, MolX’s sparse autoencoder (SAE) module also provides fine-grained interpretability at the small-molecule level. Each SAE dictionary feature corresponds to a reusable activation pattern that can be localised not only to protein residues but also to individual ligand atoms. To quantify this relationship, we compute the mean activation value of each dictionary feature across all atoms within a molecule, and visualise these values using colour intensity, where darker colours indicate stronger activation.

As shown in Fig. 6b, we selected the ten best-predicted small molecules from the PDBbind test set and visualised their atomic activation patterns. Despite structural diversity across these ligands, high-activation regions consistently localise to chemically meaningful substructures, such as aromatic rings, heterocycles, and polar functional groups involved in binding. Importantly, these activation patterns are not uniformly distributed across the molecule; instead, they concentrate on specific atoms and substructures, indicating that the SAE features capture localised chemical motifs rather than global molecular properties.

Fig. 6c shows that a t-SNE projection of sparse autoencoder activation features for PDBbind test-set molecules yields a clear separation according to normalised pKd, with high- and low-affinity compounds forming distinct regions, indicating that the learned activation space captures affinity-relevant relationships. Fig. 6d complements this global view with a single-molecule example (PROTAC 1409), where a heatmap of the three highest-mean activation features reveals distinct, feature-specific activation patterns across the molecule, suggesting that different dictionary features respond to different chemically or interaction-relevant substructures.

To further demonstrate the mechanistic interpretability of these activation features, we present a detailed case study on a PROTAC molecule targeting BCL2 in Fig. 7a-c and the one targeting BRD9 in Fig. 7d-e. The PROTAC is first coloured by its atom-level activation values, revealing three regions with particularly strong activation. We then employ the exmol framework to generate counterfactual molecules by randomly substituting functional groups at these high-activation sites, while keeping the remaining molecular scaffold unchanged (Fig. 7b). For each counterfactual, we evaluate the change in MolX’s prediction for the corresponding regression or classification task. This procedure allows us to directly probe how perturbations to activation-relevant substructures affect model outputs.

The results show that substitutions in highly activated regions lead to substantial and systematic changes in predicted activity, whereas modifications in low-activation regions have comparatively minor effects. Notably, counterfactual molecules with high structural similarity can still exhibit markedly different predictions, highlighting that MolX’s responses are sensitive to functionally relevant chemical motifs rather than overall molecular similarity alone.

Finally, Fig. 7c summarises the counterfactual analysis by aggregating prediction changes across functional group substitutions and mapping them to molecular descriptors. Using descriptor-level statistics, we quantify the positive or negative contribution of different functional groups to PROTAC prediction outcomes. For example, heteroatom-containing aromatic groups and amide-related motifs exhibit strong positive contributions, whereas certain carbonyl or nonpolar substitutions lead to reduced predicted activity. These trends align with known structure–activity relationships in PROTAC design and provide a chemically interpretable explanation for the model’s behaviour.

## Discussion

Accurately modelling protein-ligand interactions remains challenging because existing computational approaches either rely on sequence-level abstractions that overlook three-dimensional geometry, or incorporate structural information in a fragmented manner that fails to jointly capture pocket–ligand interaction patterns. In contrast, MolX explicitly represents both protein pockets and small molecules as three-dimensional graphs and learns their interaction-aware representations through large-scale pretraining, leading to consistent performance gains across 13 classification and regression benchmarks spanning PROTACs, molecular glues, ADCs, and conventional binding affinity tasks. These results demonstrate that jointly modelling pocket and ligand geometry within a unified foundation model is critical for robust and transferable protein-ligand property prediction.

Two core design choices contribute to the strong capability of MolX. First, the use of E(3)-equivariant graph Transformer layers enables joint modelling of protein pockets and ligands while preserving fundamental geometric symmetries, allowing the model to capture both local interactions and longer-range spatial dependencies that are critical for complex binding and induced-proximity scenarios. Second, the sparse autoencoder–based interpretability module decomposes latent representations into a compact set of activation features, facilitating attribution of predictions to specific protein regions and molecular substructures. Together, these components allow MolX to combine high predictive accuracy with mechanistic interpretability, bridging data-driven learning and structural insight.

Despite its advantages, MolX has several limitations that point to directions for future work. The reliance on high-quality three-dimensional structures may restrict applicability in settings where accurate pocket conformations are unavailable or highly dynamic. In addition, although the sparse autoencoder yields interpretable activation features, the correspondence between learned features and specific biochemical mechanisms is indirect and may benefit from further integration with experimental annotations or causal perturbation studies. Addressing these challenges may further extend the scope of MolX and enhance its utility for structure-based drug discovery.

## Methods

### Model details of MolX

In the pre-training procedure of MolX, the pockets and small molecules from the pre-training dataset are first converted into 3D graphs, and then noised by adding coordinate noise and atom-type masks. Then, the noisy 3D pocket/molecule graphs are encoded by a graph-based equivariant Transformer into dense representations. We set both supervised and unsupervised learning strategies to pre-train MolX. As for supervised training, two independent prediction heads comprising multilayer perceptrons are adopted to predict the homo lumo energy gap and LogP values from the encoded features. We also want to encourage MolX to learn the 3D structural and molecule conformation information during pretraining. To this end, we design two self-supervised tasks for 3D coordinate and atom-type recovery. The main idea is to recover the correct 3D positions and atom types given the corrupted input atoms and positions. The pre-training objective is given by:

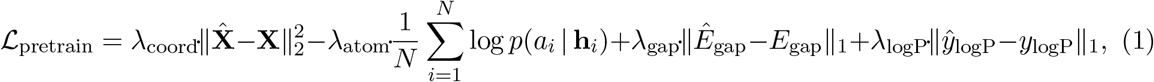

where *λ*_coord_, *λ*_atom_, *λ*_gap_ and *λ*_logP_ are balancing coefficients between different loss terms, and **R** and **R**^^^ are ground-truth and predicted node coordinates, respectively. During fine-tuning, the model is initialised from pre-training, followed by the addition of property prediction heads tailored to different downstream classification and regression tasks for further optimisation. The fine-tuning objective is given by

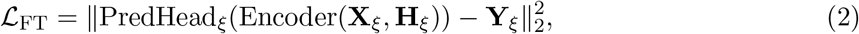

where **X***_ξ_* and **H***_ξ_* denote input coordinate matrix and feature matrix for task *ξ*, respectively. The illustration of the overall pre-training and fine-tuning procedures is exhibited in Fig. 1 (b).

As shown in Fig. 1(c), our EGNN-based Transformer follows the overall architecture of Graphormer [19], but incorporates two novel modules tailored to our pre-training dataset and self-supervised setting. Specifically, a feature unification module is introduced to integrate heterogeneous graph- and text-based molecular representations from PCQM4Mv2 and MolTextNet, and atom-type masking and coordinate-noising modules are added to facilitate self-supervised pre-training.

The model consists of an embedding layer followed by multiple Transformer-based update layers. In the embedding layer, atom types are encoded using an element dictionary that maps each type to a scalar feature. Edges are defined by chemical bonds between atoms. The node features are then encoded through stacked MolX layers. To capture potential long-range molecular interactions, we utilise Transformer layers as the backbone. Considering the intrinsic 3D spatial information of molecules, we incorporate 3D graph structural information into attention bias via an invariant 3D positional encoding module. Each MolX layer is denoted by

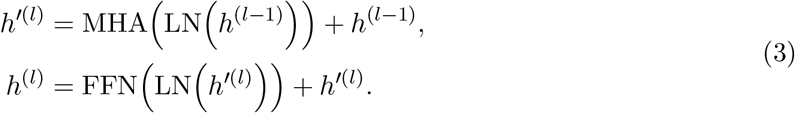

Here, layer normalisation (LN) is applied before the multi-head self-attention (MHA) and the feed-forward blocks (FFN). As shown in Fig. 1 (c), node centrality is calculated by centrality encoding, which assigns each node two real-valued embedding vectors according to its indegree and outdegree. The centrality of each node is then added to the node feature as the initial node embeddings:

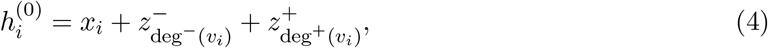

where *z^−^* and *z*^+^ *∈* R*^d^* are learnable embedding vectors specified by the indegree deg*^−^*(*v_i_*) and outdegree deg^+^(*v_i_*), respectively. As for spatial position encoding, we utilise a 3D spatial positional encoding that is invariant to global rotation and translation. Specifically, *ϕ*(*v_i_, v_j_*) : *V × V →* R denotes the Euclidean distances between atom pairs (*v_i_, v_j_*), followed by a pair-type aware Gaussian kernel. Then, the (*i, j*)-element of the Query-Key product matrix **A** is calculated as:

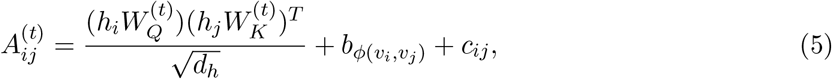

where *h_i_* denotes the representation of atom *i*, 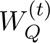 and 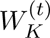 are the query and key projection matrices for the (*t*)-th attention head, and *b_ϕ_*_(_*_vi,vj_* _)_ represents the structural bias indexed by *ϕ*(*v_i_, v_j_*). *c_ij_* indicates the attention bias indexed by edge features of node pair (*v_i_, v_j_*). Specifically, for each ordered node pair (*v_i_, v_j_*), we first calculate the shortest path *SP_ij_* = (*e*_1_*, e*_2_*, …, e_N_* ) from *v_i_* to *v_j_*. edge encoding *c_ij_* is then denoted as:

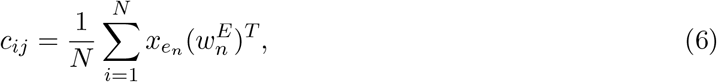

where *x_en_* is the feature of the *n*-th edge *e_n_* in *SP_ij_*, 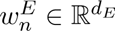 is the *n*-th weight embedding.

In addition to the basic framework of MolX, we also leverage the sparse autoencoder to find interpretable features in our foundation model. Inspired by [65, 35], we employ sparse dictionary learning to extract dictionary features from latent features of the foundation model. Let a set of vectors 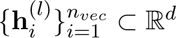 denote the internal activation features of the *l−*th EGNN-based Transformer layer in MolX, each 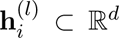 is composed of a sparse linear combination of unknown vectors 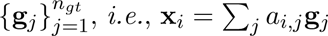 where **a***_i_* is a sparse vector. The goal of the sparse dictionary learning is to learn a dictionary of vectors, *i.e.*, dictionary features 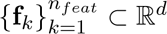 which satisfy that for any **g***_j_*, there exists a **f***_k_* such that **g***_j_ ≃* **f***_k_*.

For the input vector **h** *∈* {**h***_i_*}, our sparse autoencoder produces the output vector **x**^ by:

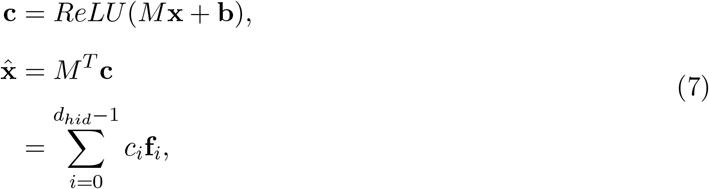

where 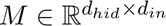 and 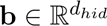 are learned parameters. *M* is the dictionary matrix, consisting of *d_hid_*rows of dictionary features **f***_i_*. For each input vector **h**, its corresponding activation feature is **c**. The sparse autoencoder is trained to minimise the following loss function:

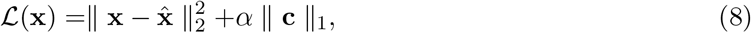

where *α* is a hyperparameter that controls the sparsity of the reconstruction. The *l*-1 loss term on *c* encourages our reconstruction to be a sparse linear combination of the dictionary features.

### Datasets

All the datasets used in pre-training and fine-tuning are listed in Table 6. We combine PCQM4Mv2 [66] and MoltextNet [38] as a 5M molecule dataset and use a 3M candidate protein pocket dataset derived from RCSB PDB [67] to pre-train a unified backbone model. Homo-lumo energy gap and LogP are used as training labels for molecules from PCAQM4Mv2 and MoltextNet, respectively. We adopt two types of downstream tasks, *i.e.*, classification and regression tasks targeting molecule-pocket-related prediction. The splitting methods are different for different tasks to maintain chemical diversity or to test out-of-distribution generalisation. All the splittings adhere to standard practices in the literature to ensure fair comparisons.

**Table 6:**
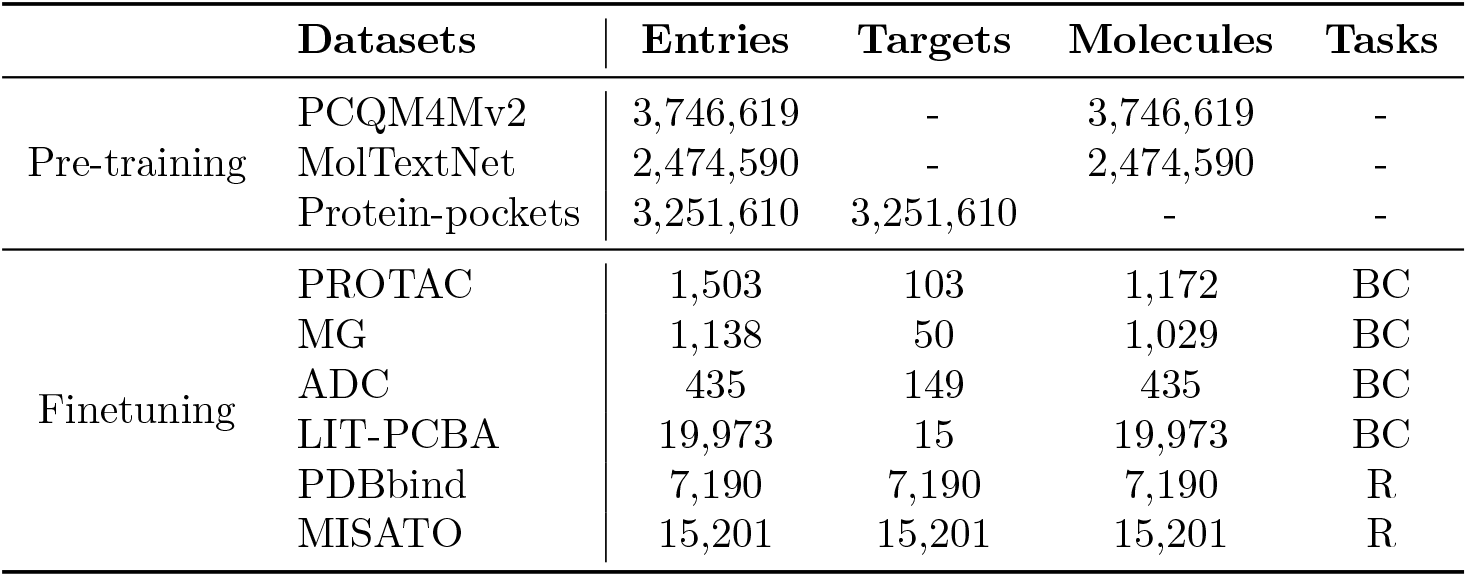
Statistics of the pretrained and finetuned datasets. BC indicates binary classification, and R indicates regression.

For the PROTAC dataset, we follow [25] and derive samples from PROTAC-DB [68]. To prevent data leakage, any test samples sharing the same target protein and PROTAC SMILES as training samples are removed [69]. MG [70] is derived from MolGlueDB [71], ADC is obtained from ADCdb [72], and Lit-PCBA is an open-source target–compound benchmark [73]. For all classification datasets, we adopt a scaffold and UMAP splits with an 80/10/10 ratio for training, validation, and test sets. PDBbind (v2020) is obtained from [45], and MISATO is obtained from [46]. For the MISATO and PDBbind-general regression tasks, we use the same scaffold- and UMAP-based splitting strategy. For PDBbind-refined, we follow a benchmark-oriented evaluation protocol and use CASF-16, OODTest, and FEP as independent external test sets.

### Training

MolX uses the EGNN base configuration (12 EGNN-based transformer layers with 32 attention heads each) with a prediction head connected to the output of the graph embeddings, consisting of a two-layer MLP with ReLU [74] activation and layer normalization [75] in between. Pretraining was carried out for 500,000 steps using a batch size of 32 for both pockets and ligands distributed across 4 A100 GPUs (making an effective batch size of 128 samples). The learning rate was linearly increased to 2 × 104 during the warm-up stage (first 60,000 steps), followed by a polynomial decaying learning rate schedule with a power of 1 and an end learning rate of 1*e −* 9. Gradient norms were clipped at 5.0, and no weight decay was used.

For fine-tuning, only the weights of the prediction head were randomly initialised. Models were trained for 100 epochs using a batch size of 32 protein-ligand pairs. The model was evaluated on the validation set every 10 epochs, and only the weights from the best-performing model according to these validation metrics were retained for further evaluation in the test set. We ran hyperparameter optimisation to find the best learning rate (1*e−*4, 2*e−*4, 5*e−*4, 1*e−*5, 1*e−*6, 1*e−*7) and dropout rate (0, 0.1, 0.15 in the prediction head) with a 5-fold cross-validation. During training, the learning rate was linearly increased during the first 16% of the training steps, followed by a polynomial decaying learning rate schedule with a power of 1.

### Baselines

The baselines include pre-training methods and non-pre-training methods. In terms of pre-training molecule prediction approaches, our baselines cover current state-of-the-art methods, including MolE [6], FradNMI [76], Coord [18], Transformer-M [19], TorchMD-Net [20], EGNN [16], SphereNet [77], Atom3D [15], HoloProt [78], AEV-PLIG [48], EHIGN [79], and DrugCLIP [80]. Coord shares the same GNN backbone with FradNMI. Transformer-M is a competitive model consisting of denoising and energy prediction pre-training tasks. We also adopt the representative approaches designed for property prediction tasks without pre-training. The models comprise general E(3)-equivariant graph neural networks (EGNN and SphereNet), and structure-based methods targeting binding affinity prediction (Atom3D and HoloProt). We use the publicly available code of the original paper to produce the results for all the baseline methods, under the same dataset split as our model.

## Data availability

All the pre-training and fine-tuning data are publicly available. PCQM4Mv2 is an Open Graph Benchmark (OGB) collection at https://ogb.stanford.edu/docs/lsc/pcqm4mv2/. MoltexNet is downloaded from https://huggingface.co/datasets/liuganghuggingface/moltextnet. Protein pockets are retrieved from the RCSB PDB database at https://www.rcsb.org/. The PROTAC dataset is retrieved from the PROTAC-DB database at https://cadd.zju.edu.cn/protacdb/downloads. The MG dataset is available at https://www.molgluedb.com. The ADC dataset is constructed from ADCdb at https://adcdb.idrblab.net/. The PCBA dataset is downloaded from https://drugdesign.unistra.fr/LIT-PCBA. The PDBbind dataset is downloaded from https://www.pdbbind-plus.org.cn/download. The MISATO dataset is downloaded from https://huggingface.co/MISATO-dataset.

## Code availability

The source code of MolX is publicly available at the GitHub repository: https://github.com/ABILiLab/MolX. The pretraining and fine-tuning datasets are publicly available on the Hugging Face repository: https://huggingface.co/datasets/JieLiu2518/MolX.

## Acknowledgments

South Australian immunoGENomics Cancer Institute (SAiGENCI) received grant funding from the Australian Government. F.L. was supported by the Australian National Health and Medical Research Council (NHMRC) Investigator Fellowship (2041439).

## Author information

F.L. conceptualised and supervised the project with help from J.S.. J.L. developed and implemented the foundation model with feedback from F.L.. J.L., T.P., X.G., Z.R., and Y.H. conduct the benchmarking analysis with feedback from F.L.. Y.Y., A.P.N., and S.P. contributed to scientific discussions and conceptual refinement of the study. All authors wrote, read, and approved the final manuscript.

## Supplementary information

Supplementary Information.pdf

