## supplementary material for "MolX: A Geometric Foundation Model for Protein-Ligand Modelling"

#### 1 Evaluation Metrics

We report complementary metrics for classification, regression, and virtual-screening evaluation. For binary classification tasks, accuracy (ACC) measures the fraction of correctly classified samples:

$$\text{ACC} = \frac{TP + TN}{TP + TN + FP + FN}, \tag{1}$$

where $TP$, $TN$, $FP$, and $FN$ denote true positives, true negatives, false positives, and false negatives, respectively. The F1 score is the harmonic mean of precision and recall:

$$\text{F1} = \frac{2 \cdot \text{Precision} \cdot \text{Recall}}{\text{Precision} + \text{Recall}}, \quad \text{Precision} = \frac{TP}{TP + FP}, \quad \text{Recall} = \frac{TP}{TP + FN}. \tag{2}$$

The area under the receiver operating characteristic curve (AUROC) evaluates the ranking quality of predicted scores across all classification thresholds. Equivalently, it can be interpreted as the probability that a randomly selected positive sample receives a higher predicted score than a randomly selected negative sample:

$$\text{AUROC} = \frac{1}{N_+ N_-} \sum_{i:y_i=1} \sum_{j:y_j=0} \mathbf{1}(s_i > s_j), \tag{3}$$

where $s_i$ is the predicted score and $N_+$ and $N_-$ are the numbers of positive and negative samples.

For regression tasks, we use mean absolute error (MAE), root mean squared error (RMSE), and Spearman's rank correlation coefficient. Given ground-truth values $y_i$ and predictions $\hat{y}_i$ for $N$ samples,

$$\text{MAE} = \frac{1}{N} \sum_{i=1}^{N} |y_i - \hat{y}_i|, \quad \text{RMSE} = \sqrt{\frac{1}{N} \sum_{i=1}^{N} (y_i - \hat{y}_i)^2}. \tag{4}$$

Spearman correlation measures monotonic agreement between predicted and true rankings:

$$\rho_s = 1 - \frac{6 \sum_{i=1}^{N} d_i^2}{N(N^2 - 1)}, \tag{5}$$

where $d_i$ is the difference between the ranks of $y_i$ and $\hat{y}_i$. Lower MAE and RMSE indicate better regression accuracy, whereas higher Spearman correlation indicates better rank consistency.

For the LIT-PCBA virtual-screening benchmark, we additionally report the enrichment factor (EF) at the top $\alpha$ fraction of ranked compounds:

$$\mathrm{EF}_\alpha = \frac{h_\alpha / n_\alpha}{H/N}, \tag{6}$$

where $N$ is the total number of compounds, $H$ is the total number of active compounds, $n_\alpha = \lfloor \alpha N \rfloor$ is the number of compounds selected from the top-ranked fraction, and $h_\alpha$ is the number of actives recovered within this subset. We report EF@0.01, EF@0.05, and EF@0.10; higher values indicate stronger early enrichment of active molecules.

### 2 Baseline Algorithms

We compare MolX with a broad set of molecular, geometric, protein-ligand, and contrastive representation learning baselines. These methods cover 2D molecular representation learning, 3D equivariant modelling, atomistic neural networks, protein structure encoders, and protein-ligand interaction models.

- **MolE** [1]: a molecular representation learning baseline designed to encode compound-level chemical information for downstream molecular property prediction.
- **FradNMI** [2]: a molecular pre-training baseline that learns chemically informative representations from molecular structures using self-supervised objectives.
- **Coord** [3]: a coordinate-based baseline that directly uses atom types and 3D coordinates to model molecular geometry.
- **Transformer-M** [4]: a transformer-based molecular model that combines graph connectivity with spatial information for molecular representation learning.
- **TorchMD-Net** [5]: an atomistic neural network originally developed for molecular dynamics and 3D molecular property prediction, using geometric information from atomic coordinates.
- **EGNN** [6]: an E(n)-equivariant graph neural network that updates node features and coordinates while preserving equivariance to rotations, translations, and reflections.
- **SphereNet** [7]: a 3D geometric graph neural network that incorporates distances, angles, and torsion information through spherical message passing.
- **Atom3D** [8]: a benchmark-oriented 3D biomolecular learning baseline that represents local atomic environments for structure-based prediction tasks.
- **HoloProt** [9]: a protein structure representation method that encodes holo-protein geometric context for downstream protein-related prediction.
- **AEV-PLIG** [10]: a protein-ligand interaction baseline based on atom-centred environment vectors, capturing local chemical environments around ligand and pocket atoms.
- **EHIGN** [11]: an equivariant hierarchical interaction graph network tailored for protein-ligand binding affinity prediction by modelling intermolecular interactions.
- **DrugCLIP** [12]: a contrastive protein-ligand representation learning method that aligns drug and target representations for interaction prediction.

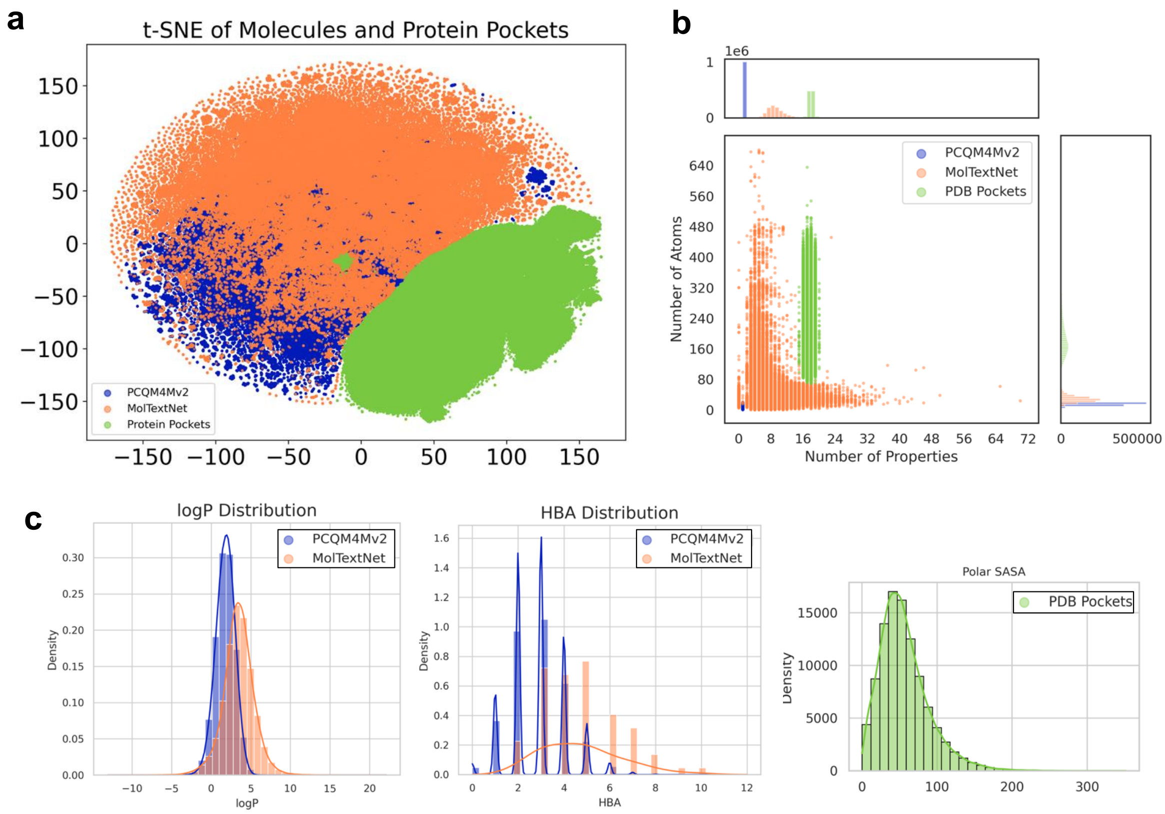


Figure S1: Statistical overview of the three pre-training datasets. a. t-SNE visualisation of molecules and protein pockets, illustrating the global distribution and overlap of PCQM4Mv2, MolTextNet, and PDB pocket datasets in the representation space. b. Dataset-level statistics, including the number of atoms and annotated properties per molecule or pocket. c. Distributions of representative physicochemical properties, including logP, hydrogen bond acceptors (HBA), and polar solvent-accessible surface area (SASA), demonstrating the complementary chemical and structural coverage of the three pre-training datasets.

### 3 Dataset statistics

All the datasets used in pre-training and fine-tuning are listed in Table S1. We combine PCQM4Mv2 and MoltextNet as a 5M-molecule dataset and use a 3M-candidate protein pocket dataset derived from RCSB PDB to pre-train a unified backbone model. Homo-lumo energy gap and LogP are used as training labels for molecules from PCAQM4Mv2 and MoltextNet, respectively. No pocket-specific property annotations were introduced, as such annotations are scarce and often task-dependent, which would limit the generality and transferability of the pre-trained model. Instead, pocket structural information and atom types are incorporated through unsupervised objectives. We adopt two types of downstream tasks, *i.e.*, classification and regression tasks targeting molecule-pocket related prediction. The splitting methods are different for different tasks to maintain chemical diversity or to test out-of-distribution generalisation. All the splittings adhere to standard practices in the literature to ensure fair comparisons.

For downstream evaluation, we adopted task-specific data splitting strategies to assess both interpolation performance and out-of-distribution generalisation. For all classification tasks, as well as the MISATO and PDBbind-general regression tasks, we used both scaffold split [13] and UMAP split [13] with an 80/20/10 ratio for training/validation/test sets. The scaffold split separates molecules by Bemis-Murcko scaffold, ensuring that compounds in the test set contain core chemical structures not shared with the training set. This setting evaluates whether a model can generalise to novel chemotypes rather than rely on memorising close analogues. The UMAP split first projects molecular or complex representations into a low-dimensional manifold and then partitions the data according to the resulting representation space. Compared with random splitting, this creates a more challenging distribution-shift setting by separating regions of the chemical or structural landscape, thereby testing whether the model can extrapolate to underrepresented or distant regions.

For PDBbind-refined, we followed a benchmark-oriented evaluation protocol using CASF-16 [14], OODTest [10], and FEP [15] as independent test sets. CASF-16 is a widely used structure-based affinity benchmark derived from high-quality protein-ligand complexes and is used to evaluate binding affinity prediction under a curated and standardised setting. OODTest is designed to measure out-of-distribution generalisation, where the test complexes differ from the training distribution in terms of target families, ligands, or binding-site environments. The FEP test set provides a more stringent evaluation of congeneric ligand series commonly used in free-energy perturbation studies, emphasising the model's ability to rank subtle affinity changes among structurally similar compounds. Together, these splits provide complementary views of model robustness: scaffold and UMAP splits assess chemical and representation-level generalisation, whereas CASF-16, OODTest, and FEP evaluate transferability to established structure-based affinity benchmarks and challenging external test scenarios.

Table S1: Statistics of the pretrained and finetuned datasets. BC indicates binary classification, and R indicates regression.

| | **Datasets** | **Entries** | **Targets** | **Molecules** | **Tasks** |
|---|---|---|---|---|---|
| Pre-training | PCQM4Mv2 | 3,746,619 | - | 3,746,619 | - |
| | MolTextNet | 2,474,590 | - | 2,474,590 | - |
| | Protein-pockets | 3,251,610 | 3,251,610 | - | - |
| Finetuning | PROTAC | 1,503 | 103 | 1,172 | BC |
| | MG | 1,138 | 50 | 1,029 | BC |
| | ADC | 435 | 149 | 435 | BC |
| | LIT-PCBA | 19,973 | 15 | 19,973 | BC |
| | PDBbind | 7,190 | 7,190 | 7,190 | R |
| | MISATO | 15,201 | 15,201 | 15,201 | R |

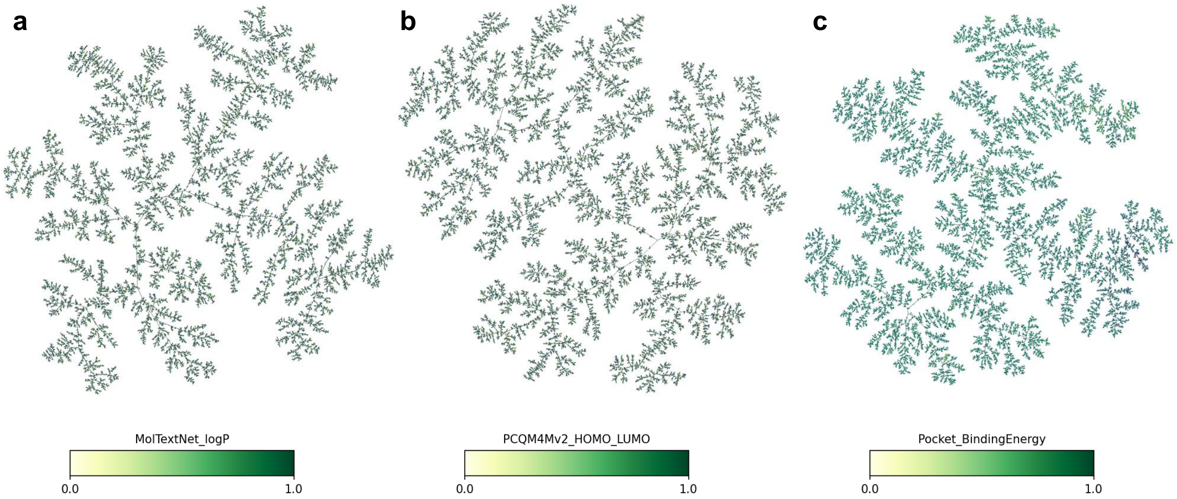


Figure S2: tmap visualisation of the three pre-trained datasets (MoltextNet, PCQM4Mv2, and Protein pocket), where edges in the figure represent local neighbourhood relationships derived from the minimum spanning tree of the k-nearest-neighbour graph in the learned representation space. a. MoltextNet visualisation, nodes are colored by logP values. b. PCQM4Mv2 visualisation, nodes are colored by homo-lumo values. c. Pocket visualisation, nodes are colored by binding energy.

### 4 Pretraining-downstream overlap analysis

To assess potential data leakage between the MolX pretraining corpus and the downstream evaluation benchmarks, we performed an exact structural overlap analysis between the pretraining inputs and the test sets used in downstream evaluation. Ligands were canonicalised with RDKit, and exact ligand overlap was computed between the unique canonical ligands in the pretraining corpus and those in each downstream test set. Protein/pocket overlap was assessed by exact sequence matching when sequence information was available.

The pretraining corpus contained 5,546,773 unique canonical ligands, with 39 invalid or unparsable ligand strings. Under the sequence extraction procedure used in this analysis, no unique protein/pocket sequences were recovered from the pretraining pocket LMDB file; therefore, protein/pocket overlap results should be interpreted as no detectable exact sequence overlap under the current parsing procedure rather than as definitive evidence that no protein-level overlap exists.

Table S2: Exact overlap between MolX pretraining inputs and downstream test sets. Ligand overlap was computed using RDKit-canonicalized ligand strings. Protein/pocket overlap was computed by exact sequence matching where sequence information was available.

| Dataset | Unique test ligands | Invalid ligands | Ligand overlap | Ligand overlap rate | Unique test protein/pocket seqs. | Protein/pocket overlap |
|---|---|---|---|---|---|---|
| PROTAC | 107 | 0 | 0 | 0.00% | 47 | 0 |
| LIT-PCBA | 1659 | 0 | 36 | 2.17% | 14 | 0 |
| ADC | 14 | 0 | 0 | 0.00% | 0 | 0 |
| Molecular glue | 100 | 0 | 0 | 0.00% | 26 | 0 |
| PDBbind-general | 395 | 29 | 12 | 3.04% | 434 | 0 |

Overall, no exact ligand overlap was observed for the PROTAC, molecular glue, or ADC test sets. Limited exact ligand overlap was detected for LIT-PCBA and PDBbind-general, with 36 overlapping ligands among 1,659 unique LIT-PCBA test ligands (2.17%) and 12 overlapping ligands among 395 unique PDBbind-general test ligands (3.04%). No exact protein/pocket sequence overlap was detected for any downstream benchmark under the current sequence-matching procedure.

Importantly, this analysis evaluates overlap in structural inputs rather than downstream label leakage. MolX pretraining uses ligand and pocket data for representation learning and does not use downstream binding affinity, activity, degradation, molecular glue, or ADC labels. Therefore, the limited ligand overlap observed in LIT-PCBA and PDBbind-general does not indicate direct label leakage. Nevertheless, we report these overlaps explicitly because self-supervised pretraining may still benefit from repeated exposure to ligand or pocket structures, and such overlap should be considered when interpreting downstream generalisation performance.

### 5 Classification results

In this section, we provide additional details on comparing model performance for classification tasks under both UMAP-based and random-based data splits.

#### 5.1 Classification results under Umap split

Table S3 shows the model performance for classification under the UMAP split. As LIT-PCBA is also widely used as a virtual screening benchmark, we additionally evaluate all models using the enrichment factor (EF). The EF results on LIT-PCBA are summarised in Table S4.

Table S3: Classification tasks under UMAP split. The best results on each dataset are in bold, the second-to-the-best results are underlined.

| **Models** | **ADC** | | | **PROTAC** | | | **MG** | | | **LIT-PCBA** | | |
|---|---|---|---|---|---|---|---|---|---|---|---|---|
| | ACC | AUC | F1 | ACC | AUC | F1 | ACC | AUC | F1 | ACC | AUC | F1 |
| MolE | **0.8627** | 0.8954 | 0.8589 | 0.8116 | 0.8521 | 0.8069 | 0.9576 | 0.9177 | 0.9569 | 0.7392 | 0.8026 | 0.7592 |
| FradNMI | 0.7450 | 0.6867 | 0.8059 | 0.6778 | 0.7516 | 0.3684 | 0.9339 | 0.9152 | 0.9514 | 0.5048 | 0.5112 | 0.5134 |
| Coord | 0.6667 | 0.6365 | 0.7301 | 0.6623 | 0.6821 | 0.3953 | 0.9406 | 0.9406 | 0.9651 | 0.5396 | 0.6024 | 0.6676 |
| Transformer-M | 0.6086 | 0.7152 | 0.3665 | 0.6233 | 0.6357 | 0.4787 | 0.9152 | 0.7980 | 0.8960 | 0.6012 | 0.6257 | 0.5969 |
| TorchMD-Net | 0.6078 | 0.8395 | 0.5077 | 0.6233 | 0.8572 | 0.621 | 0.8644 | 0.7611 | 0.5629 | 0.6482 | 0.7250 | 0.6359 |
| EGNN | 0.5294 | 0.4351 | 0.3461 | 0.6233 | 0.4473 | 0.3840 | 0.8728 | 0.7288 | 0.4660 | 0.5269 | 0.5329 | 0.5226 |
| SphereNet | 0.7647 | 0.7098 | 0.7483 | 0.6233 | 0.6754 | 0.3840 | 0.8728 | 0.7294 | 0.5245 | 0.6061 | 0.6562 | 0.6059 |
| Atom3D | 0.5294 | 0.4606 | 0.6923 | 0.7662 | 0.8361 | 0.6842 | 0.9830 | 0.9954 | 0.9901 | 0.6687 | 0.7297 | 0.6705 |
| HoloProt | 0.5294 | 0.3302 | 0.6923 | 0.5750 | 0.5639 | 0 | **0.9915** | 0.9968 | **0.9940** | 0.4965 | 0.5436 | 0 |
| AEV-PLIG | 0.2941 | 0.3388 | 0.3999 | 0.8831 | 0.6574 | 0.2500 | 0.8389 | 0.9689 | 0.8994 | 0.4897 | 0.4980 | 0.5366 |
| EHIGN | 0.3529 | 0.5345 | 0.2667 | 0.8961 | 0.6844 | 0.1999 | 0.9756 | **0.9970** | 0.9851 | 0.6449 | 0.7054 | 0.6252 |
| DrugCLIP | 0.8375 | 0.8708 | 0.8500 | 0.6338 | 0.6425 | 0 | 0.8474 | 0.7508 | 0.9150 | 0.5611 | 0.5698 | 0.5762 |
| MolX | 0.8465 | **0.9115** | **0.8721** | **0.9110** | **0.8893** | **0.8359** | 0.9801 | 0.9755 | 0.9851 | **0.7596** | **0.8281** | **0.7633** |

Table S4: Enrichment factor (EF) results on LIT-PCBA under UMAP split. The best results on each dataset are in bold, the second-to-the-best results are underlined.

| **Model** | **EF@0.01** | **EF@0.05** | **EF@0.10** |
|---|---|---|---|
| MolE | 1.9863 | 1.9092 | 1.8313 |
| FradNMI | 1.3904 | 1.1684 | 1.0710 |
| Coord | 1.6884 | 1.3242 | 1.2950 |
| Transformer-M | 1.7026 | 1.2728 | 1.2790 |
| TorchMD-Net | 1.6885 | 1.7142 | 1.6467 |
| EGNN | 0.8940 | 0.9542 | 1.1006 |
| SphereNet | 1.6886 | 1.5003 | 1.3920 |
| Atom3D | 1.8918 | 1.7935 | 1.6763 |
| HoloProt | 1.4188 | 1.0992 | 1.2112 |
| AEV-PLIG | 0.7567 | 0.9642 | 0.9689 |
| EHIGN | 1.6628 | 1.6385 | 1.5417 |
| DrugCLIP | 1.3904 | 1.2853 | 1.1879 |
| MolX | **1.9931** | **1.9315** | **1.8804** |

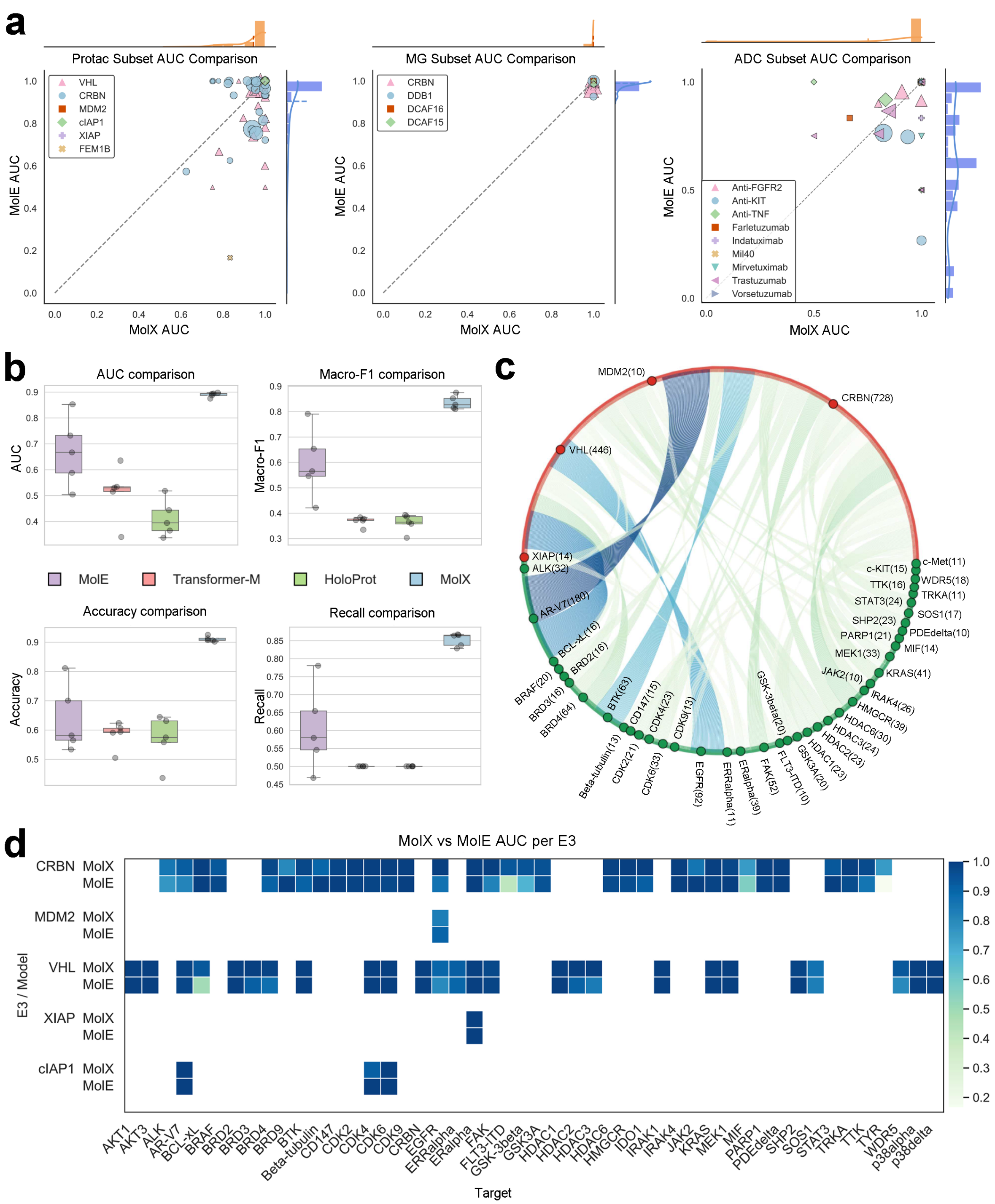

Figure S3: **Performance evaluation results on classification tasks under UMAP split.** **a**. Subset-level AUC comparison across classification tasks. **b**. Classification performance across multiple metrics. **c**. Fine-grained decomposition of the PROTAC dataset by target–E3 pairs. **d**. Heat map comparison of MolX and MolE across PROTAC target–E3 subsets.

#### 5.2 Classification Results under Random Split

Table S5 shows the model performance for classification under a random split. As LIT-PCBA is also widely used as a virtual screening benchmark, we additionally evaluate all models using the enrichment factor (EF). The EF results on LIT-PCBA are summarised in Table S6.

Table S5: Model performance on downstream classification tasks under random split. The best results on each dataset are in bold, the second-to-the-best results are underlined.

| Models | ADC | | | PROTAC | | | MG | | | LIT-PCBA | | |
|---|---|---|---|---|---|---|---|---|---|---|---|---|
| | ACC | AUC | F1 | ACC | AUC | F1 | ACC | AUC | F1 | ACC | AUC | F1 |
| MolE | 0.833 | 0.884 | 0.762 | 0.725 | 0.700 | 0.793 | 0.876 | 0.914 | 0.834 | 0.716 | 0.788 | 0.716 |
| FradNMI | 0.738 | 0.806 | 0.671 | 0.699 | 0.732 | 0.649 | 0.774 | 0.500 | 0.436 | 0.544 | 0.566 | 0.534 |
| Coord | 0.651 | 0.667 | 0.631 | 0.576 | 0.592 | 0.319 | 0.719 | 0.847 | 0.495 | 0.519 | 0.525 | 0.386 |
| Transformer-M | 0.643 | 0.619 | 0.524 | 0.596 | 0.381 | 0.537 | 0.726 | 0.547 | 0.450 | 0.518 | 0.534 | 0.481 |
| TorchMD-Net | 0.571 | 0.554 | 0.640 | 0.733 | 0.842 | 0.535 | 0.797 | 0.733 | 0.873 | 0.663 | 0.732 | 0.650 |
| EGNN | 0.667 | 0.573 | 0.4 | 0.620 | 0.527 | 0.383 | 0.761 | 0.600 | 0.432 | 0.542 | 0.551 | 0.542 |
| SphereNet | 0.667 | 0.722 | 0.800 | 0.620 | 0.695 | 0.637 | 0.761 | 0.705 | 0.857 | 0.623 | 0.681 | 0.612 |
| Atom3D | 0.609 | 0.634 | 0.596 | 0.609 | 0.395 | 0.378 | 0.948 | 0.994 | 0.931 | 0.666 | 0.728 | 0.666 |
| HoloProt | 0.636 | 0.758 | 0.389 | 0.580 | 0.605 | 0.367 | 0.965 | 0.989 | 0.957 | 0.506 | 0.511 | 0.350 |
| AEV-PLIG | 0.813 | 0.580 | 0.885 | 0.695 | 0.646 | 0.233 | 0.9210 | 0.966 | 0.941 | 0.502 | 0.531 | 0.348 |
| EHIGN | 0.604 | 0.529 | 0.738 | 0.880 | 0.849 | 0.181 | 0.956 | 0.996 | 0.965 | 0.645 | 0.709 | 0.700 |
| DrugCLIP | 0.857 | 0.833 | 0.909 | 0.654 | 0.697 | 0.000 | 0.740 | 0.854 | 0.834 | 0.548 | 0.554 | 0.532 |
| MolX | **0.9318** | **0.9807** | **0.9538** | **0.8433** | **0.9211** | **0.8365** | **0.9668** | **0.9972** | **0.9767** | **0.7526** | **0.8135** | **0.7633** |

Table S6: Enrichment factor (EF) results on LIT-PCBA under Random split. The best results on each dataset are in bold, the second-to-the-best results are underlined.

| Model | EF@0.01 | EF@0.05 | EF@0.10 |
|---|---|---|---|
| MolE | 2.0181 | 1.9778 | 1.9273 |
| FradNMI | 1.6456 | 1.3685 | 1.2045 |
| Coord | 1.4736 | 1.1515 | 1.1859 |
| Transformer-M | 0.8072 | 1.2109 | 1.2008 |
| TorchMD-Net | 1.8600 | 1.8250 | 1.6581 |
| EGNN | 1.7575 | 1.2103 | 1.1052 |
| SphereNet | 1.4470 | 1.2501 | 1.2436 |
| Atom3D | 2.0000 | 1.8600 | 1.7900 |
| HoloProt | 0.9000 | 1.1800 | 1.1100 |
| AEV-PLIG | 0.8000 | 0.9800 | 1.0800 |
| EHIGN | 1.8784 | 1.6203 | 1.5219 |
| DrugCLIP | 1.2124 | 1.1658 | 1.2243 |
| MolX | **2.2590** | **2.1362** | **1.9615** |

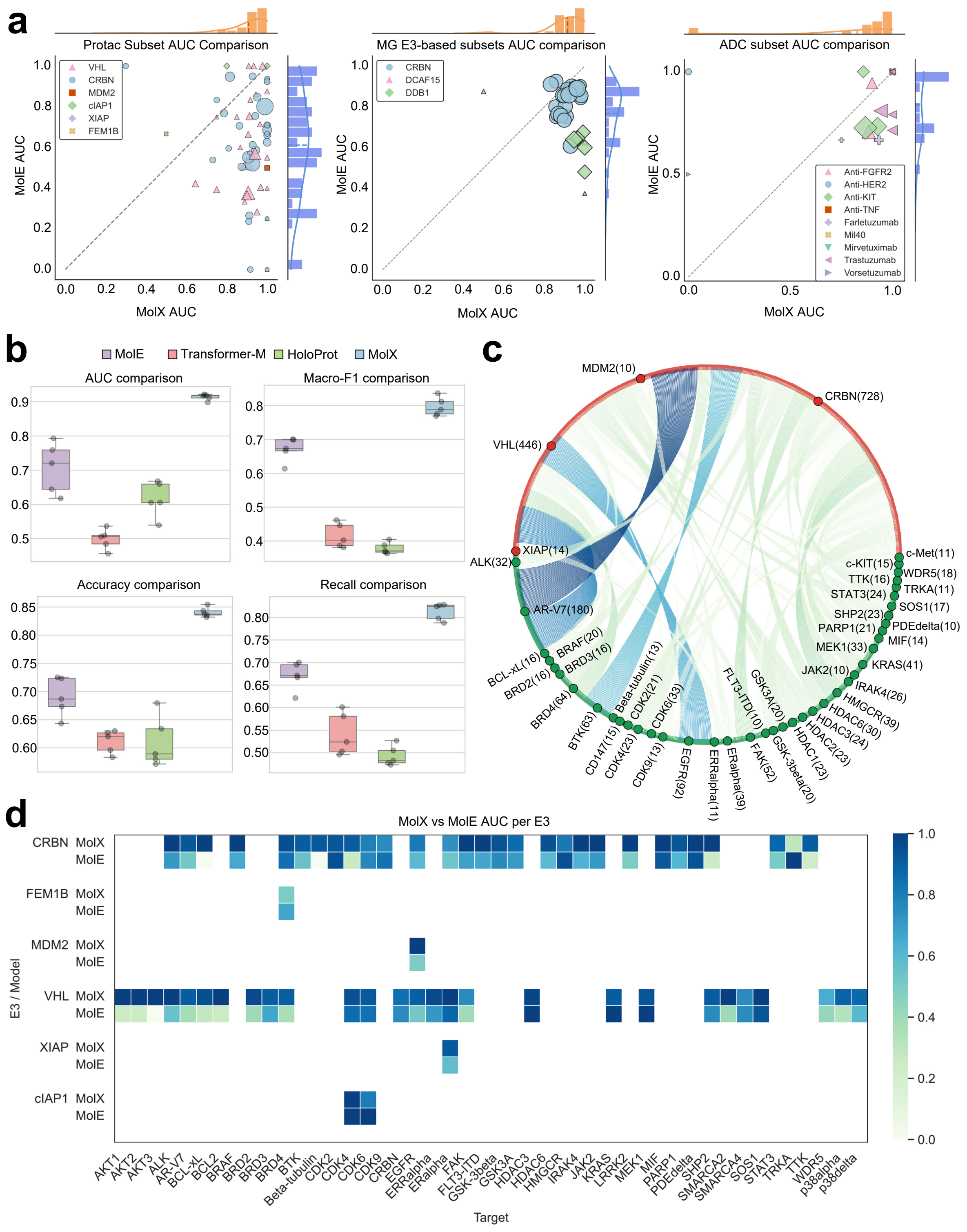


Figure S4: **Performance evaluation results on classification tasks under random split.** **a**. Subset-level AUC comparison across classification tasks. **b**. Classification performance across multiple metrics. **c**. Fine-grained decomposition of the PROTAC dataset by target–E3 pairs. **d**. Heat map comparison of MolX and MolE across PROTAC target–E3 subsets.

### 6 Regression results

In this section, we provide additional details on comparing model performance for regression tasks under both UMAP-based and random-based data splits.

#### 6.1 Regression results under UMAP split

Table S7: Model performance on PDBbind dataset under UMAP split. The best results on each dataset are in bold, the second-best results are underlined.

| Models | Kd | | | Ki | | | IC50 | | |
|---|---|---|---|---|---|---|---|---|---|
| | MAE | RMSE | Spearman | MAE | RMSE | Spearman | MAE | RMSE | Spearman |
| MolE | 1.2399 | 1.5823 | 0.4794 | 1.3726 | 1.6638 | 0.6116 | <u>1.0987</u> | <u>1.3972</u> | **0.6913** |
| FradNMI | 1.7338 | 2.3520 | 0.3244 | 1.9126 | 2.6653 | 0.3370 | 1.6751 | 2.1539 | 0.3381 |
| Coord | 1.5170 | 1.9495 | 0.3694 | 1.5588 | 1.8940 | 0.4885 | 1.3800 | 1.7723 | 0.4242 |
| Transformer-M | 1.3940 | 1.7301 | 0.4192 | 1.4627 | 1.7994 | 0.4647 | 1.3625 | 1.7061 | 0.4727 |
| TorchMD-Net | **1.1841** | <u>1.4816</u> | <u>0.5308</u> | 1.2887 | 1.5910 | 0.6288 | 1.2023 | 1.5139 | 0.5787 |
| EGNN | 1.3211 | 1.6554 | 0.3916 | 1.4527 | 1.7953 | 0.4846 | 1.3557 | 1.6923 | 0.3778 |
| SphereNet | 1.3118 | 1.6316 | 0.3738 | 1.4477 | 1.7757 | 0.5005 | 1.3676 | 1.6904 | 0.3991 |
| Atom3D | 1.4654 | 1.8264 | 0.2676 | 1.5538 | 1.9037 | 0.4246 | 1.2935 | 1.5833 | 0.4065 |
| HoloProt | 1.4786 | 1.8216 | 0.3304 | 1.5413 | 1.9236 | 0.4413 | 1.3372 | 1.6282 | 0.5098 |
| AEV-PLIG | 1.2675 | 1.6001 | 0.4147 | 1.4012 | 1.7006 | 0.6433 | 1.2410 | 1.5081 | 0.6433 |
| EHIGN | 1.2150 | 1.5296 | 0.4901 | <u>1.1829</u> | <u>1.4750</u> | 0.6453 | 1.1594 | 1.4663 | 0.6839 |
| DrugCLIP | 1.2616 | 1.5807 | 0.4283 | 1.3714 | 1.6608 | <u>0.6469</u> | 1.1411 | 1.4167 | 0.5573 |
| MolX | <u>1.2028</u> | **1.4359** | **0.6337** | **1.1669** | **1.4675** | **0.6919** | **1.0649** | **1.3796** | <u>0.6840</u> |

Table S8: Model performance on MISATO dataset under UMAP split, EA, EN, $\eta$, IP, KPT and MW is the abbreviation for Electron Affinity, Electronegativity, Hardness, Ionisation Potential, Koopman value, and Molecular Weight, respectively.

| Models | EA | | EN | | $\eta$ | | IP | | KPT | | MW | |
|---|---|---|---|---|---|---|---|---|---|---|---|---|
| | MAE | RMSE | MAE | RMSE | MAE | RMSE | MAE | RMSE | MAE | RMSE | MAE | RMSE |
| FradNMI | 0.4023 | 0.5274 | 0.4033 | 0.5245 | 0.3121 | 0.4872 | 0.4375 | 0.5612 | 0.4158 | 0.5357 | 0.9549 | 1.2631 |
| Coord | 0.3811 | 0.5090 | 0.4129 | 0.5425 | 0.2959 | 0.4510 | 0.4477 | 0.5947 | 0.4059 | 0.5487 | 1.0190 | 1.3099 |
| TorchMD-Net | <u>0.1644</u> | <u>0.2285</u> | 0.0905 | 0.1348 | <u>0.1862</u> | <u>0.3676</u> | 0.0955 | 0.1373 | 0.1826 | 0.2843 | 0.3710 | 0.4567 |
| EGNN | 0.2615 | 0.3419 | <u>0.0588</u> | **0.1111** | 0.2346 | 0.4032 | 0.0452 | **0.0997** | <u>0.0818</u> | <u>0.2117</u> | **0.2626** | **0.3431** |
| SphereNet | <u>0.1644</u> | <u>0.2285</u> | 0.0905 | 0.1348 | <u>0.1862</u> | <u>0.3676</u> | 0.0955 | 0.1373 | 0.1826 | 0.2843 | 0.3710 | 0.4567 |
| Atom3D | 0.1759 | 0.3239 | 0.3585 | 0.3727 | 1.1509 | 1.2283 | 0.5715 | 0.5807 | 0.4373 | 0.4611 | 4.2001 | 4.2304 |
| AEV-PLIG | 2.1007 | 9.8108 | 3.6216 | 4.0030 | 5.0875 | 5.3912 | 3.4166 | 3.8159 | 3.5695 | 3.9520 | 1.3000 | 1.8279 |
| EHIGN | 17.6452 | 35.8074 | 17.7981 | 35.6565 | 18.3772 | 36.1127 | 17.7200 | 35.5899 | 17.7640 | 35.6264 | 17.2505 | 34.6603 |
| DrugCLIP | 0.1664 | 0.3380 | 0.0648 | <u>0.1191</u> | 0.2466 | 0.4301 | <u>0.0446</u> | <u>0.1004</u> | 0.0835 | 0.2143 | 0.2822 | 0.3648 |
| MolX | **0.1412** | **0.2046** | **0.0585** | 0.1287 | **0.1767** | **0.3526** | **0.0439** | 0.1066 | **0.0809** | **0.2018** | <u>0.2686</u> | <u>0.3498</u> |

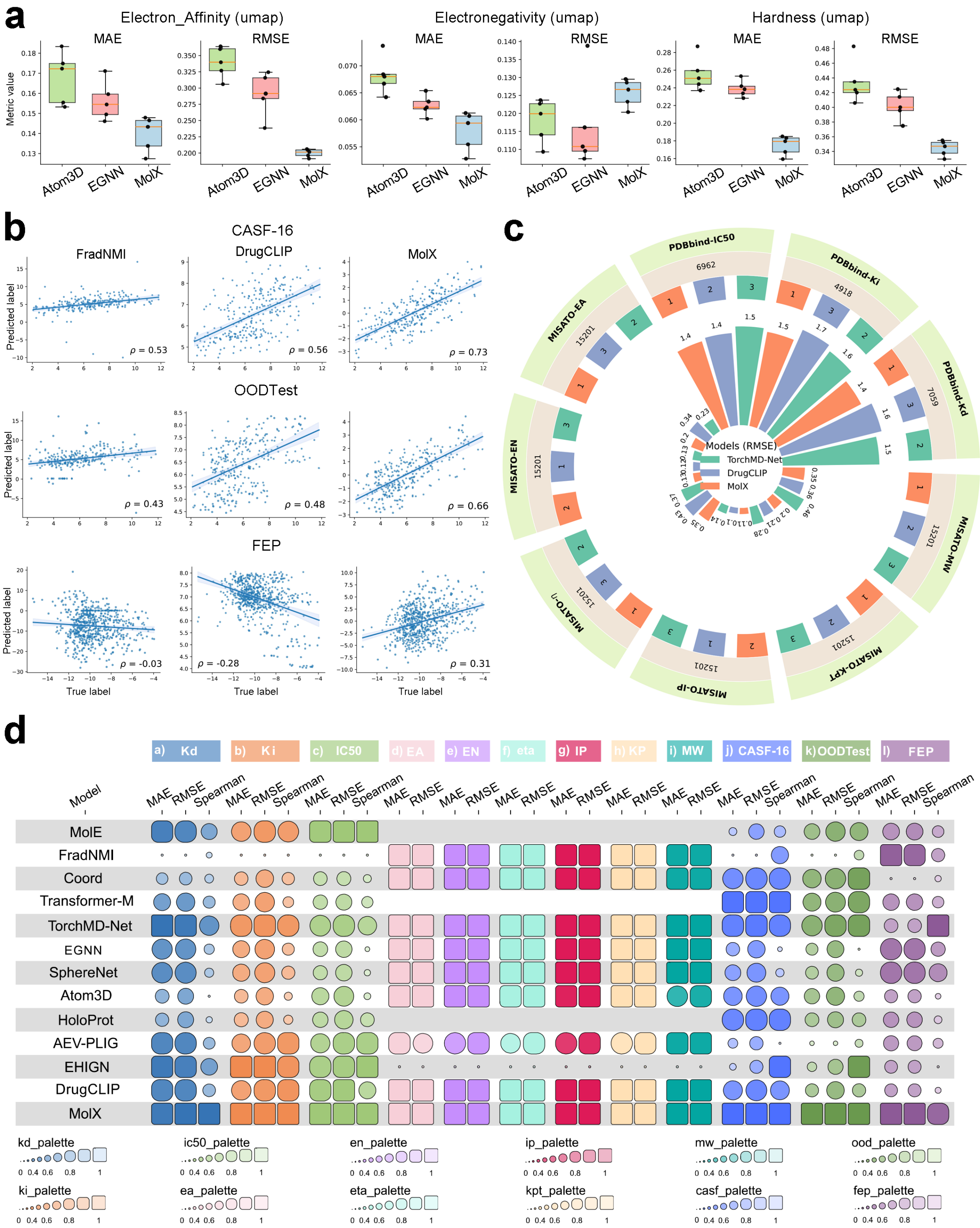


Figure S5: **Regression results under the UMAP split.** **a**. Comparison of MAE, RMSE, and Spearman correlation for predictions of electron affinity and binding affinities across representative models. **b**. Radial bar plot comparing RMSE across multiple regression tasks, including PDBbind and MISATO datasets. **c**. Scatter plots of predicted versus ground-truth values for regression of pdbbind CASF-16 and OODTest, comparing FradNMI, Atom3D, and MolX. **d**. Sensitivity analysis showing prediction trends across binned target ranges for multiple regression properties. **e**. Summary heatmap comparing regression performance (MAE, RMSE, and Spearman) of all models across pdbbind and MISATO. Larger rectangles indicate better relative performance for each metric and task.

#### 6.2 Regression results under random split

Table S9: Model performance on PDBbind dataset for regression under random split. The best results on each dataset are in bold, the second-best results are underlined.

| Models | Kd | | | Ki | | | IC50 | | |
|---|---|---|---|---|---|---|---|---|---|
| | MAE | RMSE | Spearman | MAE | RMSE | Spearman | MAE | RMSE | Spearman |
| MolE | 1.2406 | <u>1.5504</u> | 0.0893 | 1.5605 | 1.9180 | 0.1653 | 1.3377 | 1.6556 | 0.0725 |
| FradNMI | 1.7742 | 2.3445 | 0.3301 | 1.8463 | 2.4992 | 0.3748 | 1.6813 | 2.1987 | 0.4023 |
| Coord | 1.5843 | 2.0498 | 0.3326 | 1.6528 | 2.1227 | 0.5673 | 1.3395 | 1.7507 | 0.4937 |
| Transformer-M | 1.3292 | 1.7427 | 0.0994 | 1.5911 | 1.9999 | 0.0721 | 1.3336 | 1.6715 | 0.0551 |
| TorchMD-Net | **1.2078** | 1.5881 | <u>0.4320</u> | 1.4301 | 1.7369 | 0.6185 | 1.1229 | 1.4048 | 0.5557 |
| EGNN | 1.3235 | 1.7524 | 0.2305 | 1.6133 | 1.9647 | 0.4819 | 1.2784 | 1.5821 | 0.3917 |
| SphereNet | 1.2890 | 1.6636 | 0.3649 | 1.5328 | 1.8715 | 0.5044 | 1.2978 | 1.5779 | 0.3184 |
| Atom3D | 1.4773 | 1.8416 | 0.0150 | 1.7278 | 2.1132 | 0.0457 | 1.4398 | 1.7519 | 0.0033 |
| HoloProt | 1.2782 | 1.5868 | 0.3802 | <u>1.2730</u> | <u>1.6605</u> | <u>0.6305</u> | <u>1.1105</u> | <u>1.3968</u> | **0.5662** |
| MolX | <u>1.2266</u> | **1.5043** | **0.5350** | **1.2569** | **1.6530** | **0.6753** | **1.0766** | **1.3792** | <u>0.5589</u> |

Table S10: Model performance on MISATO dataset for regression under random split, EA, EN, $\eta$, IP, KPT and MW is the abbreviation for Electron Affinity, Electronegativity, Hardness, Ionisation Potential, Koopman value, and Molecular Weight, respectively.

| Models | EA | | EN | | $\eta$ | | IP | | KPT | | MW | |
|---|---|---|---|---|---|---|---|---|---|---|---|---|
| | MAE | RMSE | MAE | RMSE | MAE | RMSE | MAE | RMSE | MAE | RMSE | MAE | RMSE |
| FradNMI | 0.4175 | 0.5222 | 0.4045 | 0.4907 | 0.3017 | 0.4468 | 0.4617 | 0.5631 | 0.4254 | 0.5239 | 0.8770 | 1.1013 |
| Coord | 0.4021 | 0.5069 | 0.4265 | 0.5416 | 0.2740 | 0.4154 | 0.4689 | 0.6002 | 0.4494 | 0.5892 | 1.0350 | 1.3480 |
| TorchMD-Net | 0.2684 | 0.3408 | 0.0870 | 0.1254 | 0.2110 | 0.3914 | 0.1013 | 0.1657 | 0.1196 | 0.2055 | **0.2148** | **0.2830** |
| EGNN | 0.2391 | 0.3143 | <u>0.0610</u> | **0.1051** | 0.2422 | 0.4342 | <u>0.0461</u> | 0.1388 | <u>0.0809</u> | 0.1944 | 0.2687 | 0.3513 |
| SphereNet | 0.1567 | <u>0.2159</u> | 0.1936 | 0.2237 | <u>0.2051</u> | <u>0.3859</u> | 0.0679 | 0.1457 | 0.1097 | 0.2243 | 0.6565 | 0.7216 |
| Atom3D | <u>0.1488</u> | 0.2906 | 0.0955 | 0.1337 | 0.2323 | 0.3874 | 0.0831 | **0.1102** | 0.0922 | <u>0.1767</u> | <u>0.2420</u> | <u>0.3220</u> |
| MolX | **0.1256** | **0.1920** | **0.0607** | <u>0.1227</u> | **0.1956** | **0.3710** | **0.0454** | <u>0.1171</u> | **0.0800** | **0.1668** | 0.2507 | 0.3394 |

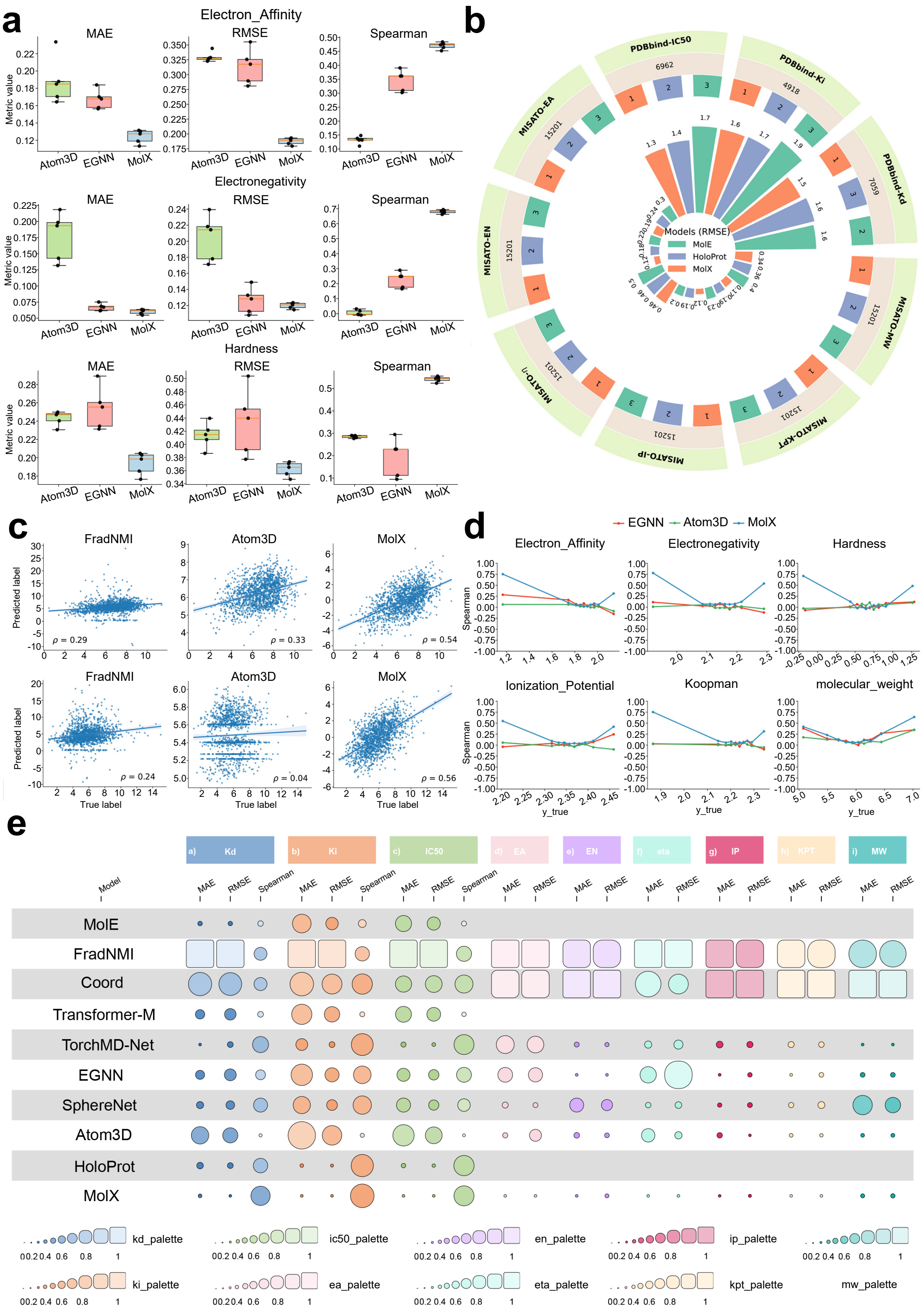


Figure S6: **Regression results under the random split.** **a**. Comparison of MAE, RMSE, and Spearman correlation for predictions of electron affinity and binding affinities across representative models. **b**. Radial bar plot comparing RMSE across multiple regression tasks, including PDBbind and MISATO datasets. **c**. Scatter plots of predicted versus ground-truth values for regression of pdbbind CASF-16 and OODTest, comparing FradNMI, Atom3D, and MolX. **d**. Sensitivity analysis showing prediction trends across binned target ranges for multiple regression properties. **e**. Summary heatmap comparing regression performance (MAE, RMSE, and Spearman) of all models across pdbbind and MISATO. Larger rectangles indicate better relative performance for each metric and task.

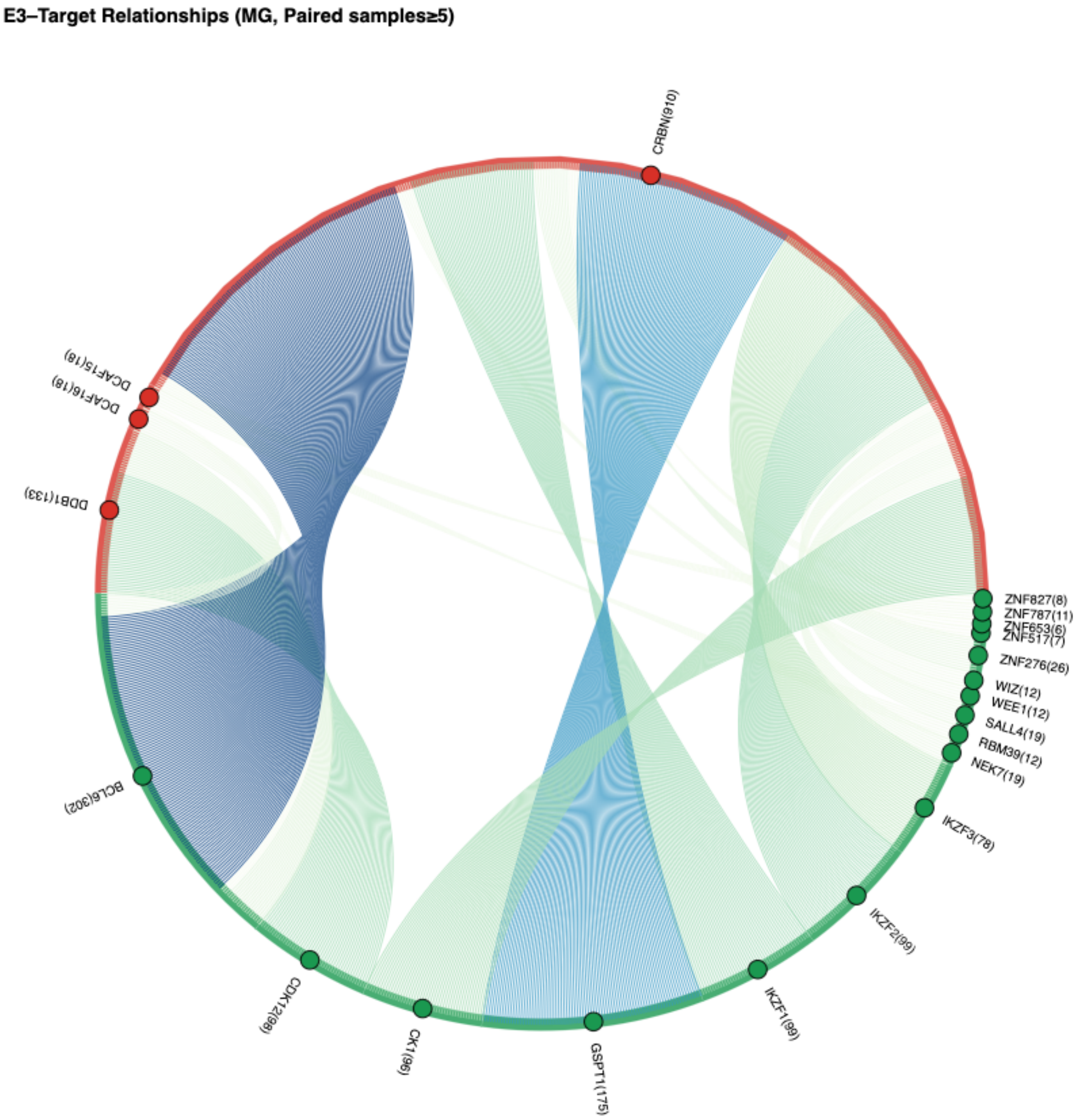


Figure S7: Fine-grained decomposition of the molecular glue dataset by target–E3 pairs.

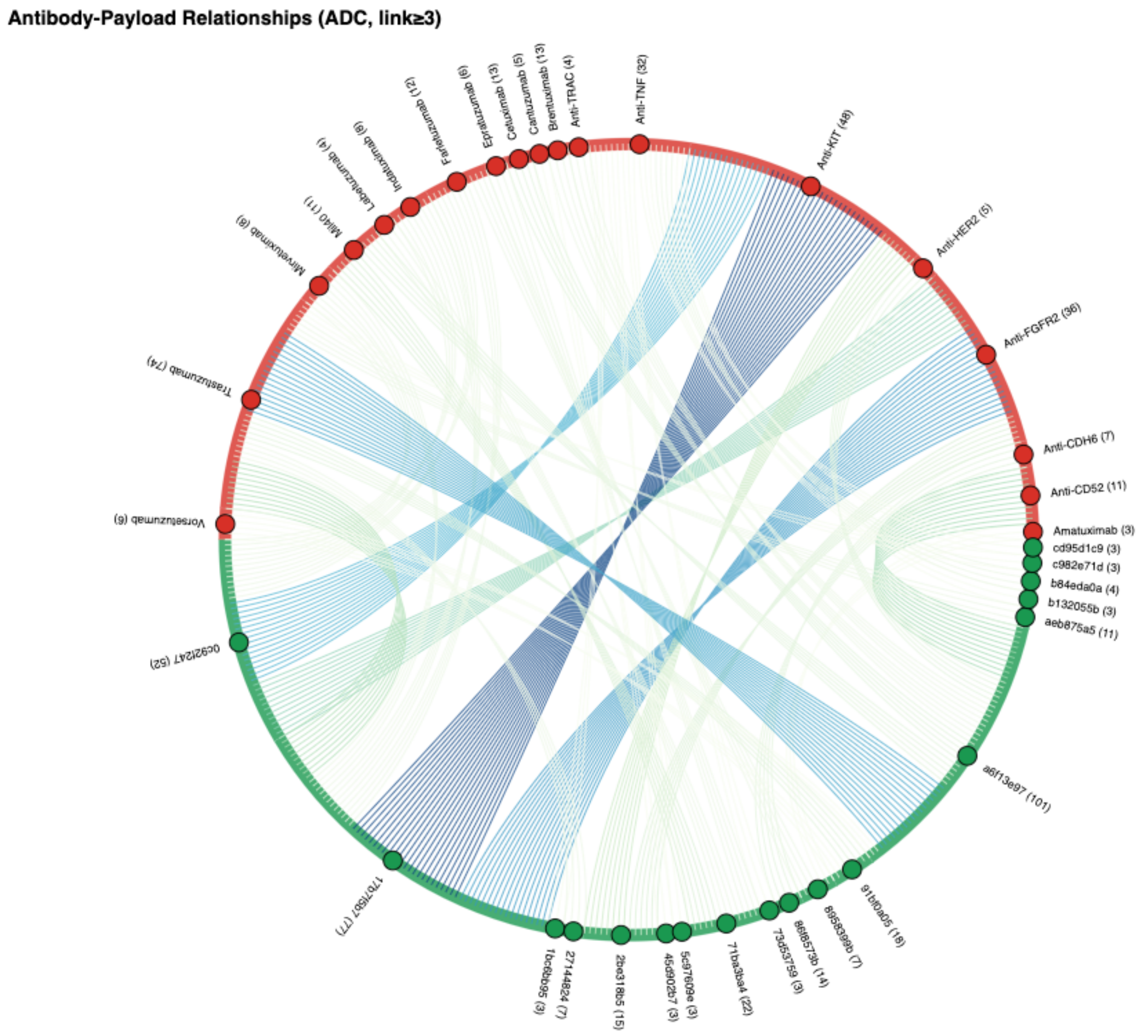


Figure S8: Fine-grained decomposition of the ADC dataset by antibody–payload pairs.

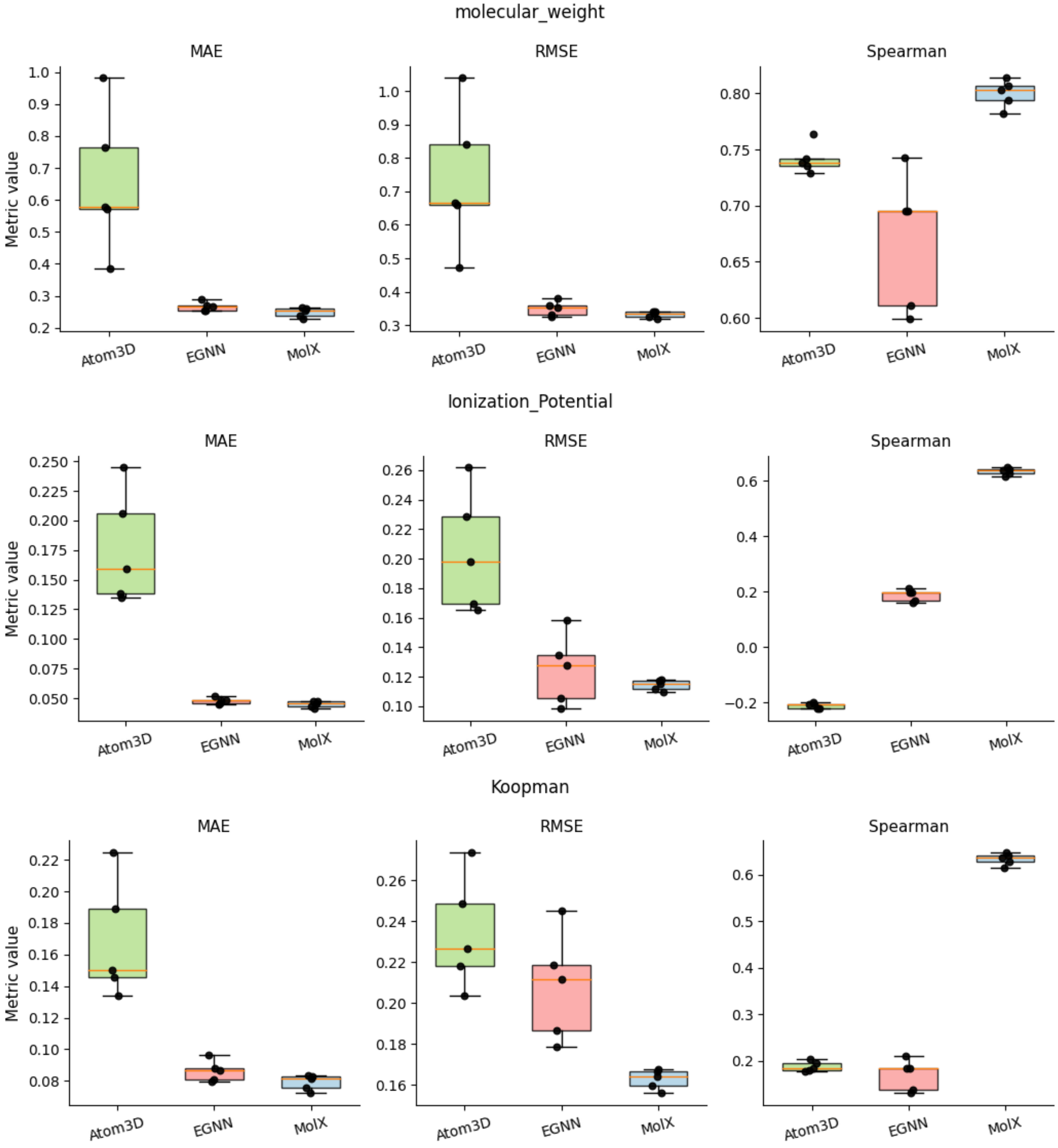


Figure S9: Comparison of MAE, RMSE, and Spearman correlation for molecular weight, Ionisation potential and Koopman predictions on the MISATO dataset across representative models.

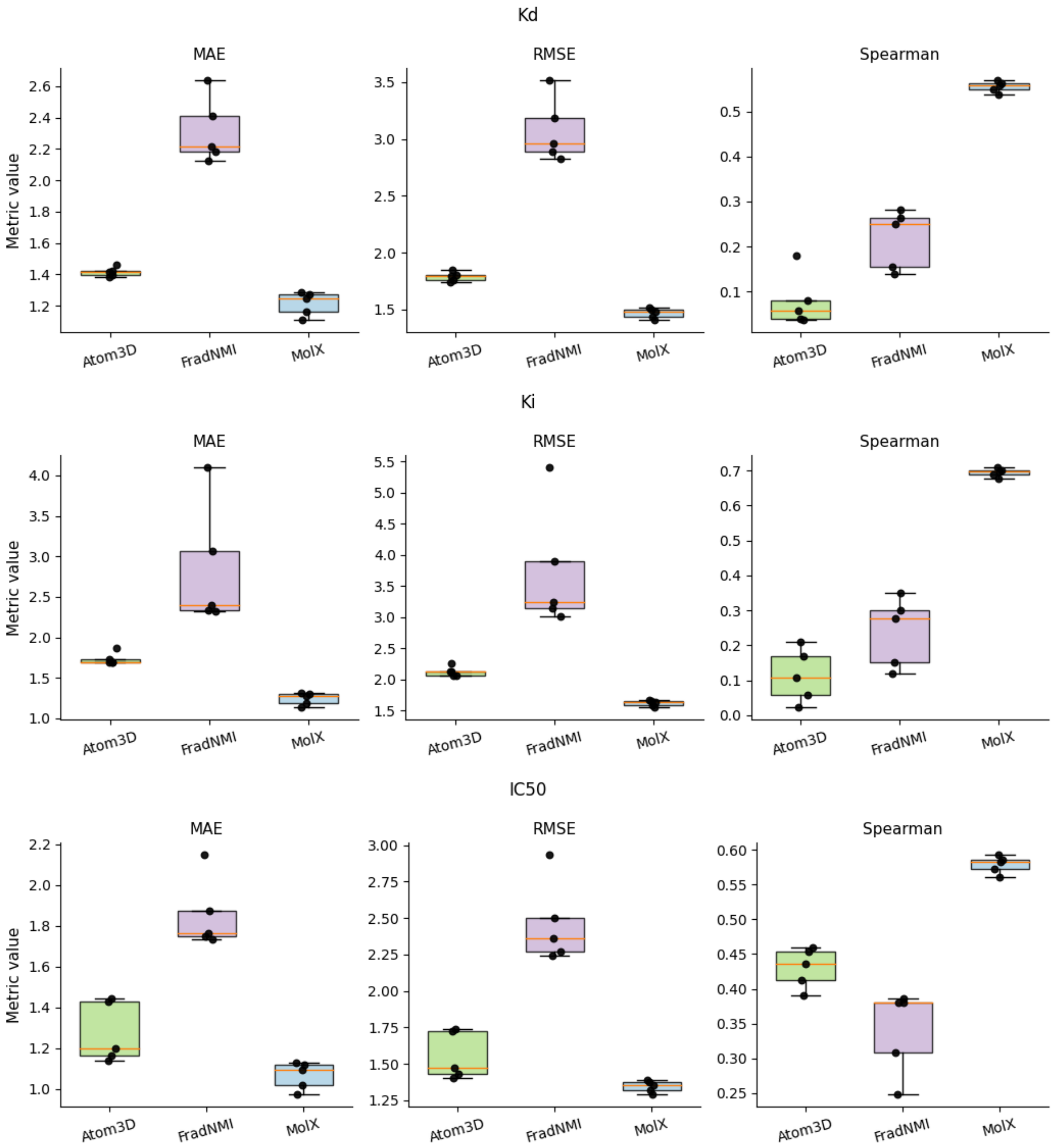


Figure S10: Comparison of MAE, RMSE, and Spearman correlation for $Kd$, $Ki$ and $IC_{50}$ predictions on PDBbind dataset across representative models

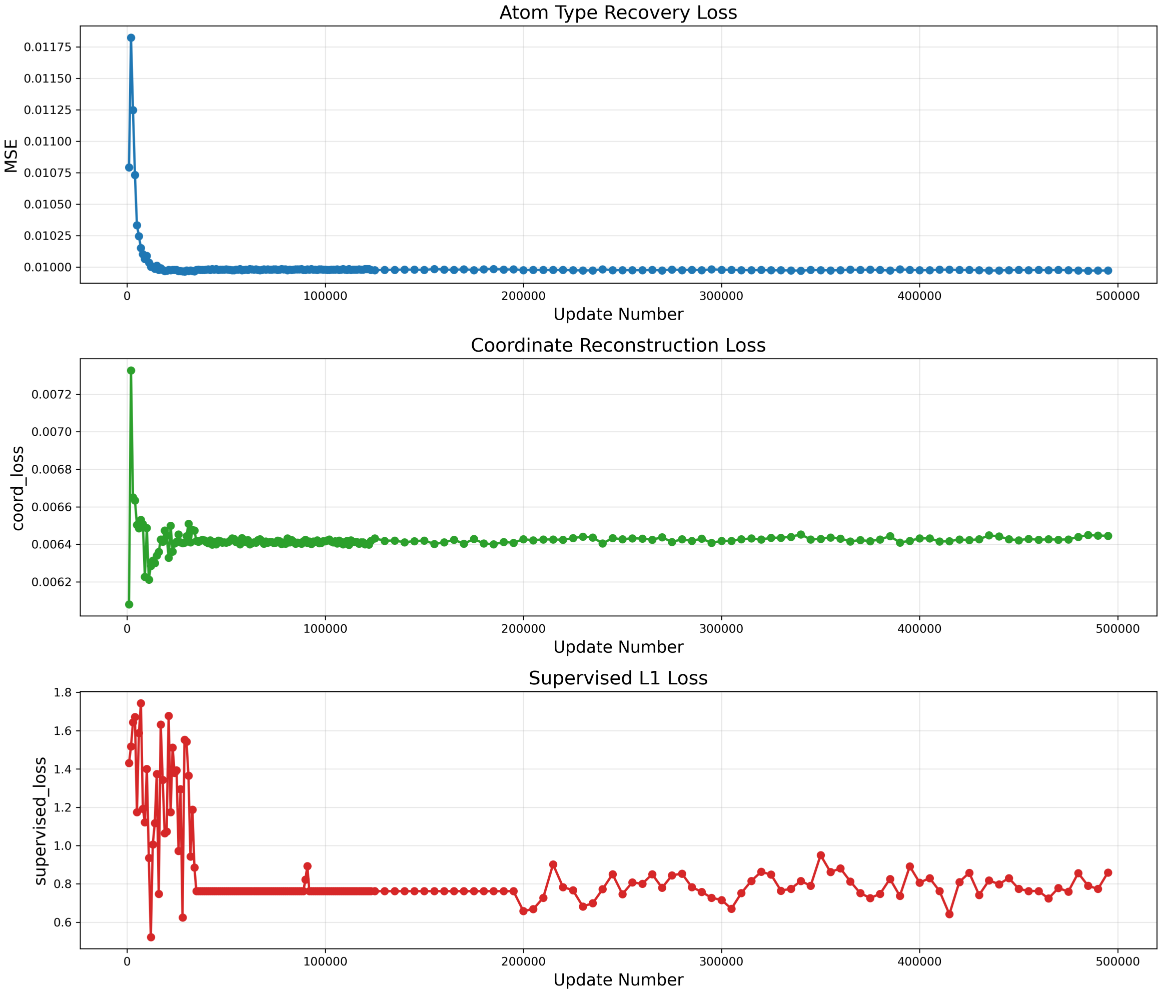


Figure S11: Evolution of pre-training and supervised losses over training steps. The atom-type recovery loss (top) and coordinate reconstruction loss (middle) converge rapidly and remain stable throughout training, demonstrating successful learning of atomic identity and geometric reconstruction. In contrast, the supervised L1 loss (bottom) shows higher variance and slower stabilisation, consistent with the increased complexity of supervised target prediction.

Table S11: Statistics relationships of activation features and E3 protein descriptors. Activation idx denotes the index of the activation feature. The target descriptor indicates the normalised functional or structural descriptor of the target protein. Count is the number of occurrences of a given activation–descriptor pair. The within-act ratio represents the fraction of this descriptor among all occurrences associated with the same activation feature. Rank in act indicates the rank of the descriptor within a given activation feature when sorted by occurrence frequency. The act total denotes the total number of descriptor occurrences associated with that activation feature.

| Activation idx | E3 descriptor | Count | Within act ratio | Rank in act | Act total |
|---|---:|---:|---:|---:|---:|
| 1075 | von Hippel-Lindau disease tumor suppressor | 27 | 0.675000 | 1 | 40 |
| 1075 | Involved in binding to CCT complex | 10 | 0.250000 | 2 | 40 |
| 1075 | Interaction with Elongin BC complex | 3 | 0.075000 | 3 | 40 |
| 1289 | Protein cereblon | 437 | 0.292308 | 1 | 1495 |
| 1289 | Lon N-terminal | 398 | 0.266221 | 2 | 1495 |
| 1289 | von Hippel-Lindau disease tumor suppressor | 313 | 0.209365 | 3 | 1495 |
| 1289 | Involved in binding to CCT complex | 146 | 0.097659 | 4 | 1495 |
| 1289 | Baculoviral IAP repeat-containing protein 2 | 49 | 0.032776 | 5 | 1495 |
| 1289 | Interaction with Elongin BC complex | 43 | 0.028763 | 6 | 1495 |
| 1289 | E3 ubiquitin-protein ligase Mdm2 | 31 | 0.020736 | 7 | 1495 |
| 1289 | ARF-binding | 20 | 0.013378 | 8 | 1495 |
| 1289 | Region II | 20 | 0.013378 | 9 | 1495 |
| 1289 | Interaction with MTBP | 15 | 0.010033 | 10 | 1495 |
| 1452 | von Hippel-Lindau disease tumor suppressor | 14 | 0.518519 | 1 | 27 |
| 1452 | Involved in binding to CCT complex | 8 | 0.296296 | 2 | 27 |
| 1452 | Protein cereblon | 4 | 0.148148 | 3 | 27 |
| 1452 | Lon N-terminal | 1 | 0.037037 | 4 | 27 |
| 1750 | von Hippel-Lindau disease tumor suppressor | 10 | 0.714286 | 1 | 14 |
| 1750 | Interaction with Elongin BC complex | 2 | 0.142857 | 2 | 14 |
| 1750 | Involved in binding to CCT complex | 2 | 0.142857 | 3 | 14 |
| 1961 | von Hippel-Lindau disease tumor suppressor | 2 | 1.000000 | 1 | 2 |
| 2061 | von Hippel-Lindau disease tumor suppressor | 23 | 0.547619 | 1 | 42 |
| 2061 | Involved in binding to CCT complex | 17 | 0.404762 | 2 | 42 |
| 2061 | Interaction with Elongin BC complex | 2 | 0.047619 | 3 | 42 |
| 211 | Involved in binding to CCT complex | 4 | 0.666667 | 1 | 6 |
| 211 | von Hippel-Lindau disease tumor suppressor | 2 | 0.333333 | 2 | 6 |
| 3035 | von Hippel-Lindau disease tumor suppressor | 4 | 0.571429 | 1 | 7 |
| 3035 | Interaction with Elongin BC complex | 2 | 0.285714 | 2 | 7 |
| 3035 | Involved in binding to CCT complex | 1 | 0.142857 | 3 | 7 |
| 3245 | Lon N-terminal | 13 | 0.464286 | 1 | 28 |
| 3245 | Protein cereblon | 8 | 0.285714 | 2 | 28 |
| 3245 | E3 ubiquitin-protein ligase Mdm2 | 2 | 0.071429 | 3 | 28 |
| 3245 | Disordered | 1 | 0.035714 | 4 | 28 |
| 3245 | Interaction with MTBP | 1 | 0.035714 | 5 | 28 |
| 3245 | Involved in binding to CCT complex | 1 | 0.035714 | 6 | 28 |
| 3245 | Necessary for interaction with USP2 | 1 | 0.035714 | 7 | 28 |
| 3245 | von Hippel-Lindau disease tumor suppressor | 1 | 0.035714 | 8 | 28 |
| 3303 | von Hippel-Lindau disease tumor suppressor | 17 | 0.500000 | 1 | 34 |
| 3303 | Involved in binding to CCT complex | 12 | 0.352941 | 2 | 34 |
| 3303 | Lon N-terminal | 5 | 0.147059 | 3 | 34 |
| 3707 | von Hippel-Lindau disease tumor suppressor | 1 | 1.000000 | 1 | 1 |
| 3801 | Involved in binding to CCT complex | 10 | 0.500000 | 1 | 20 |
| 3801 | von Hippel-Lindau disease tumor suppressor | 9 | 0.450000 | 2 | 20 |
| 3801 | Protein cereblon | 1 | 0.050000 | 3 | 20 |
| 3893 | Protein cereblon | 291 | 0.296939 | 1 | 980 |
| 3893 | Lon N-terminal | 265 | 0.270408 | 2 | 980 |
| 3893 | von Hippel-Lindau disease tumor suppressor | 174 | 0.177551 | 3 | 980 |
| 3893 | Involved in binding to CCT complex | 123 | 0.125510 | 4 | 980 |
| 3893 | Interaction with Elongin BC complex | 30 | 0.030612 | 5 | 980 |
| 3893 | Baculoviral IAP repeat-containing protein 2 | 24 | 0.024490 | 6 | 980 |
| 3893 | E3 ubiquitin-protein ligase Mdm2 | 16 | 0.016327 | 7 | 980 |
| 3893 | ARF-binding | 15 | 0.015306 | 8 | 980 |
| 3893 | Disordered | 15 | 0.015306 | 9 | 980 |
| 3893 | Interaction with MTBP | 14 | 0.014286 | 10 | 980 |

Table S12: Statistical relationships of activation features and E3 protein descriptors for additional activation features not included in Table S1.

| Activation idx | E3 descriptor | Count | Within act ratio | Rank in act | Act total |
|---|---|---|---|---|---|
| 516 | von Hippel-Lindau disease tumor suppressor | 17 | 0.320755 | 1 | 53 |
| 516 | Protein cereblon | 13 | 0.245283 | 2 | 53 |
| 516 | Lon N-terminal | 10 | 0.188679 | 3 | 53 |
| 516 | Involved in binding to CCT complex | 9 | 0.169811 | 4 | 53 |
| 516 | ARF-binding | 1 | 0.018868 | 5 | 53 |
| 516 | E3 ubiquitin-protein ligase Mdm2 | 1 | 0.018868 | 6 | 53 |
| 516 | Necessary for interaction with USP2 | 1 | 0.018868 | 7 | 53 |
| 516 | Region II | 1 | 0.018868 | 8 | 53 |

Table S13: Statistical relationships of activation features and target protein descriptors.

| Activation idx | Target descriptor | Count | Within act ratio | Rank in act | Act total |
|---|---|---|---|---|---|
| 1075 | Androgen receptor | 6 | 0.222222 | 1 | 27 |
| 1075 | Disordered | 5 | 0.185185 | 2 | 27 |
| 1075 | Interaction with LPXN | 5 | 0.185185 | 3 | 27 |
| 1075 | Interaction with ZNF318 | 4 | 0.148148 | 4 | 27 |
| 1075 | Interaction with KAT7 | 3 | 0.111111 | 5 | 27 |
| 1075 | Interaction with CCAR1 | 2 | 0.074074 | 6 | 27 |
| 1075 | NR LBD | 2 | 0.074074 | 7 | 27 |
| 1289 | Protein kinase | 58 | 0.194631 | 1 | 298 |
| 1289 | Androgen receptor | 28 | 0.093960 | 2 | 298 |
| 1289 | Dual specificity mitogen-activated protein kinase | 24 | 0.080537 | 3 | 298 |
| 1289 | Interaction with LPXN | 21 | 0.070470 | 4 | 298 |
| 1289 | NR LBD | 20 | 0.067114 | 5 | 298 |
| 1289 | Interaction with KAT7 | 17 | 0.057047 | 6 | 298 |
| 1289 | Modulating | 15 | 0.050336 | 7 | 298 |
| 1289 | Cyclin-dependent kinase 4 | 12 | 0.040268 | 8 | 298 |
| 1289 | Interaction with CCAR1 | 12 | 0.040268 | 9 | 298 |
| 1289 | Interaction with ZNF318 | 12 | 0.040268 | 10 | 298 |
| 1452 | Protein kinase | 32 | 0.137931 | 1 | 232 |
| 1452 | Cyclin-dependent kinase 4 | 21 | 0.090517 | 2 | 232 |
| 1452 | 3-hydroxy-3-methylglutaryl-coenzyme A reductase | 18 | 0.077586 | 3 | 232 |
| 1452 | Androgen receptor | 18 | 0.077586 | 4 | 232 |
| 1452 | Interaction with LPXN | 16 | 0.068966 | 5 | 232 |
| 1452 | Tyrosine-protein kinase BTK | 16 | 0.068966 | 6 | 232 |
| 1452 | ATP | 14 | 0.060345 | 7 | 232 |
| 1452 | Interaction with CCAR1 | 11 | 0.047414 | 8 | 232 |
| 1452 | SH2 | 11 | 0.047414 | 9 | 232 |
| 1452 | Interaction with KAT7 | 10 | 0.043103 | 10 | 232 |
| 1750 | Protein kinase | 25 | 0.135135 | 1 | 185 |
| 1750 | Androgen receptor | 18 | 0.097297 | 2 | 185 |
| 1750 | Disordered | 17 | 0.091892 | 3 | 185 |
| 1750 | NR LBD | 15 | 0.081081 | 4 | 185 |
| 1750 | Interaction with KAT7 | 14 | 0.075676 | 5 | 185 |
| 1750 | Cyclin-dependent kinase 4 | 12 | 0.064865 | 6 | 185 |
| 1750 | Interaction with CCAR1 | 12 | 0.064865 | 7 | 185 |
| 1750 | Bromodomain-containing protein 4 | 11 | 0.059459 | 8 | 185 |
| 1750 | Epidermal growth factor receptor | 10 | 0.054054 | 9 | 185 |
| 1750 | Interaction with LPXN | 8 | 0.043243 | 10 | 185 |
| 1961 | Protein kinase | 71 | 0.253571 | 1 | 280 |
| 1961 | SH2 | 19 | 0.067857 | 2 | 280 |
| 1961 | Androgen receptor | 18 | 0.064286 | 3 | 280 |
| 1961 | Interaction with LPXN | 17 | 0.060714 | 4 | 280 |
| 1961 | Cyclin-dependent kinase 4 | 16 | 0.057143 | 5 | 280 |
| 1961 | Cyclin-dependent kinase 6 | 13 | 0.046429 | 6 | 280 |
| 1961 | Tyrosine-protein kinase BTK | 13 | 0.046429 | 7 | 280 |
| 1961 | ATP | 12 | 0.042857 | 8 | 280 |
| 1961 | NR LBD | 12 | 0.042857 | 9 | 280 |
| 1961 | 3-hydroxy-3-methylglutaryl-coenzyme A reductase | 11 | 0.039286 | 10 | 280 |
| 2061 | Protein kinase | 27 | 0.178808 | 1 | 151 |
| 2061 | Androgen receptor | 12 | 0.079470 | 2 | 151 |
| 2061 | NR LBD | 12 | 0.079470 | 3 | 151 |
| 2061 | 3-hydroxy-3-methylglutaryl-coenzyme A reductase | 9 | 0.059603 | 4 | 151 |
| 2061 | Interaction with KAT7 | 9 | 0.059603 | 5 | 151 |
| 2061 | Interaction with LPXN | 9 | 0.059603 | 6 | 151 |
| 2061 | ATP | 8 | 0.052980 | 7 | 151 |
| 2061 | Cyclin-dependent kinase 6 | 8 | 0.052980 | 8 | 151 |
| 2061 | Signal transducer and activator of transcription 3 | 8 | 0.052980 | 9 | 151 |
| 2061 | Bromodomain-containing protein 4 | 7 | 0.046358 | 10 | 151 |

Table S14: Statistical relationships of activation features and target protein descriptors for additional activation features not included in Table S3.

| Activation idx | Target descriptor | Count | Within act ratio | Rank in act | Act total |
|---|---|---|---|---|---|
| 211 | Protein kinase | 50 | 0.175439 | 1 | 285 |
| 211 | Cyclin-dependent kinase 6 | 27 | 0.094737 | 2 | 285 |
| 211 | Epidermal growth factor receptor | 23 | 0.080702 | 3 | 285 |
| 211 | Androgen receptor | 20 | 0.070175 | 4 | 285 |
| 211 | Interaction with KAT7 | 19 | 0.066667 | 5 | 285 |
| 211 | NR LBD | 16 | 0.056140 | 6 | 285 |
| 211 | Cyclin-dependent kinase 4 | 15 | 0.052632 | 7 | 285 |
| 211 | Interaction with CCAR1 | 15 | 0.052632 | 8 | 285 |
| 211 | 3-hydroxy-3-methylglutaryl-coenzyme A reductase | 14 | 0.049123 | 9 | 285 |
| 211 | Dual specificity mitogen-activated protein kinase | 12 | 0.042105 | 10 | 285 |
| 3035 | Interaction with CCAR1 | 6 | 0.133333 | 1 | 45 |
| 3035 | 3-hydroxy-3-methylglutaryl-coenzyme A reductase | 5 | 0.111111 | 2 | 45 |
| 3035 | Androgen receptor | 5 | 0.111111 | 3 | 45 |
| 3035 | NR LBD | 5 | 0.111111 | 4 | 45 |
| 3035 | Cyclin-dependent kinase 6 | 4 | 0.088889 | 5 | 45 |
| 3035 | Epidermal growth factor receptor | 4 | 0.088889 | 6 | 45 |
| 3035 | Interaction with LPXN | 4 | 0.088889 | 7 | 45 |
| 3035 | Signal transducer and activator of transcription 3 | 3 | 0.066667 | 8 | 45 |
| 3035 | Bromo 1 | 2 | 0.044444 | 9 | 45 |
| 3035 | Interaction with KAT7 | 2 | 0.044444 | 10 | 45 |
| 3245 | Androgen receptor | 13 | 0.106557 | 1 | 122 |
| 3245 | Protein kinase | 12 | 0.098361 | 2 | 122 |
| 3245 | Interaction with CCAR1 | 11 | 0.090164 | 3 | 122 |
| 3245 | SH2 | 9 | 0.073770 | 4 | 122 |
| 3245 | NR LBD | 8 | 0.065574 | 5 | 122 |
| 3245 | Dual specificity mitogen-activated protein kinase | 7 | 0.057377 | 6 | 122 |
| 3245 | Interaction with KAT7 | 7 | 0.057377 | 7 | 122 |
| 3245 | Interaction with LPXN | 7 | 0.057377 | 8 | 122 |
| 3245 | Bromodomain-containing protein 4 | 6 | 0.049180 | 9 | 122 |
| 3245 | Cyclin-dependent kinase 4 | 6 | 0.049180 | 10 | 122 |
| 3303 | Interaction with KAT7 | 5 | 0.192308 | 1 | 26 |
| 3303 | Interaction with LPXN | 5 | 0.192308 | 2 | 26 |
| 3303 | NR LBD | 5 | 0.192308 | 3 | 26 |
| 3303 | 3-hydroxy-3-methylglutaryl-coenzyme A reductase | 2 | 0.076923 | 4 | 26 |
| 3303 | Androgen receptor | 2 | 0.076923 | 5 | 26 |
| 3303 | Protein kinase | 2 | 0.076923 | 6 | 26 |
| 3303 | Disordered | 1 | 0.038462 | 7 | 26 |
| 3303 | Epidermal growth factor receptor | 1 | 0.038462 | 8 | 26 |
| 3303 | Interaction with CCAR1 | 1 | 0.038462 | 9 | 26 |
| 3303 | Interaction with ZNF318 | 1 | 0.038462 | 10 | 26 |
| 3707 | Androgen receptor | 2 | 0.222222 | 1 | 9 |
| 3707 | Interaction with KAT7 | 2 | 0.222222 | 2 | 9 |
| 3707 | NR LBD | 2 | 0.222222 | 3 | 9 |
| 3707 | Interaction with CCAR1 | 1 | 0.111111 | 4 | 9 |
| 3707 | Interaction with LPXN | 1 | 0.111111 | 5 | 9 |
| 3707 | Modulating | 1 | 0.111111 | 6 | 9 |
| 3801 | Disordered | 5 | 0.454545 | 1 | 11 |
| 3801 | Androgen receptor | 1 | 0.090909 | 2 | 11 |
| 3801 | Interaction with CCAR1 | 1 | 0.090909 | 3 | 11 |
| 3801 | Interaction with KAT7 | 1 | 0.090909 | 4 | 11 |
| 3801 | Interaction with LPXN | 1 | 0.090909 | 5 | 11 |
| 3801 | Modulating | 1 | 0.090909 | 6 | 11 |
| 3801 | NR LBD | 1 | 0.090909 | 7 | 11 |

Table S15: Statistical relationships of activation features and target protein descriptors for additional activation features not included in Table S3.

| Activation idx | Target descriptor | Count | Within act ratio | Rank in act | Act total |
|---|---|---|---|---|---|
| 3893 | Androgen receptor | 5 | 0.156250 | 1 | 32 |
| 3893 | Interaction with LPXN | 4 | 0.125000 | 2 | 32 |
| 3893 | Protein kinase | 4 | 0.125000 | 3 | 32 |
| 3893 | Disordered | 3 | 0.093750 | 4 | 32 |
| 3893 | 3-hydroxy-3-methylglutaryl-coenzyme A reductase | 2 | 0.062500 | 5 | 32 |
| 3893 | Bromo 1 | 2 | 0.062500 | 6 | 32 |
| 3893 | Interaction with CCAR1 | 2 | 0.062500 | 7 | 32 |
| 3893 | Interaction with KAT7 | 2 | 0.062500 | 8 | 32 |
| 3893 | NR LBD | 2 | 0.062500 | 9 | 32 |
| 3893 | Bromodomain-containing protein 4 | 1 | 0.031250 | 10 | 32 |
| 516 | Protein kinase | 41 | 0.225275 | 1 | 182 |
| 516 | Cyclin-dependent kinase 6 | 20 | 0.109890 | 2 | 182 |
| 516 | NR LBD | 12 | 0.065934 | 3 | 182 |
| 516 | Cyclin-dependent kinase 4 | 10 | 0.054945 | 4 | 182 |
| 516 | Disordered | 10 | 0.054945 | 5 | 182 |
| 516 | Tyrosine-protein kinase BTK | 10 | 0.054945 | 6 | 182 |
| 516 | Androgen receptor | 9 | 0.049451 | 7 | 182 |
| 516 | 3-hydroxy-3-methylglutaryl-coenzyme A reductase | 8 | 0.043956 | 8 | 182 |
| 516 | Bromodomain-containing protein 4 | 8 | 0.043956 | 9 | 182 |
| 516 | Interaction with CCAR1 | 8 | 0.043956 | 10 | 182 |

[3] Sheheryar Zaidi, Michael Schaarschmidt, James Martens, Hyunjik Kim, Yee Whye Teh, Alvaro Sanchez-Gonzalez, Peter Battaglia, Razvan Pascanu, and Jonathan Godwin. Pre-training via denoising for molecular property prediction. In *International Conference on Learning Representations*, 2023.

[4] Chengxuan Ying, Tianle Cai, Shengjie Luo, Shuxin Zheng, Guolin Ke, Di He, Yanming Shen, and Tie-Yan Liu. Do transformers really perform badly for graph representation? *Advances in neural information processing systems*, 34:28877–28888, 2021.

[5] Philipp Thölke and Gianni De Fabritiis. Torchmd-net: equivariant transformers for neural network based molecular potentials. *arXiv preprint arXiv:2202.02541*, 2022.

[6] Vıctor Garcia Satorras, Emiel Hoogeboom, and Max Welling. E (n) equivariant graph neural networks. In *International conference on machine learning*, pages 9323–9332. PMLR, 2021.

[7] Yi Liu, Limei Wang, Meng Liu, Xuan Zhang, Bora Oztekin, and Shuiwang Ji. Spherical message passing for 3d graph networks. *arXiv preprint arXiv:2102.05013*, 2021.

[8] Raphael John Lamarre Townshend, Martin Vögele, Patricia Adriana Suriana, Alexander Derry, Alexander Powers, Yianni Laloudakis, Sidhika Balachandar, Bowen Jing, Brandon M Anderson, Stephan Eismann, et al. Atom3d: Tasks on molecules in three dimensions. In *Thirty-fifth Conference on Neural Information Processing Systems Datasets and Benchmarks Track (Round 1)*.
